# Aminomutation catalyzed by CO_2_ self-sufficient cascade amino acid decarboxylases

**DOI:** 10.1101/2023.08.12.552924

**Authors:** Zhan Song, Yuan Li, Yunjie Li, Xiangwei Cui, Jian-Jiang Zhong, Yi-Heng P. J. Zhang

## Abstract

Molecular editing of an amino group from α-position of amino acids to its β-position is of scientific interest and could be economically appealing. Here we reconstructed an *in vitro* biotransformation pathway composed of two cascade decarboxylases, i.e., aspartate β-decarboxylase and aspartate α-decarboxylase, and implemented molecular editing to change α-alanine into β-alanine. In it, we discovered a new reaction of aspartate β-decarboxylase that can fix CO_2_ directly. This cascade enzymatic pathway enabled an aminomutation reaction with 100% carbon atom economy. This work presented the first CO_2_-fixing biological reaction catalyzed by the amino acid decarboxylases and demonstrated a new means for the molecular editing of α-amino acids.

## Introduction

Enzymatic CO_2_ fixation is one of the most important biochemical reactions for the global carbon cycle, and typical carboxylation enzymes include carboxylases and decarboxylases for their reverse reactions. Until now, the reported reversing decarboxylases for the CO_2_ fixation reactions are bivalent metal-dependent decarboxylases ^[1]^, cofactor-independent decarboxylases ^[2]^, prenylated flavin dependent decarboxylases ^[2c, 3]^, and TPP-dependent keto acid decarboxylases. ^[4]^ To our knowledge, there are no reports pertaining to the CO_2_-fixing reaction catalyzed by amino acid decarboxylases.

β-Amino acids refer to amino acids whose amino group is bound to their β carbon position. Most β-amino acids are synthesized by intramolecular rearrangement of the α-amino group at the β-position of α-amino acids. ^[5]^ β-Alanine, the most popular natural β-amino acid, is a precursor of vitamin B5 and of dipeptides such as carnosine, goose carnosine, and so on. ^[6]^ Due to its unique physiological roles, it has been widely used in the food and feed industries. ^[5, 7]^ Also, it can be used to synthesize poly-β-alanine, which is used in cosmetics, water purification, medicine, and other fields. ^[8]^

β-Alanine can be produced by chemical catalysis, microbial fermentation, and enzymatic biocatalysis. Its current global market is approximately 30,000 tonnes. Traditional method of chemical synthesis involves a reaction of malonic acid with ammonia catalyzed by sodium hydroxide or ammonium chloride. ^[9]^ However, this method suffers from the use of petrochemical resource, harsh reactions (i.e., high temperature and pressure), and salt waste generated. Microbial fermentation and enzyme biocatalysis are gradually replacing chemical catalysis. All microbial pathways are based on aspartate-α-decarboxylase (AAD, EC 4.1.1.12) that decarboxylates α-aspartate, which was synthesized from different carbon sources, such as glucose ^[10]^ and fatty acid ^[11]^. Enzymatic biocatalysis of β-alanine is of great interests due to its high chemical selectivity, modest reaction conditions, and high product yield.

Until now, there are a few reports on β-alanine synthesis by enzymatic biocatalysis. One approach is AAD-catalyzing the decarboxylation reaction of aspartate. The substrate is synthesized from fumaric acid and ammonia by aspartate ammonia-lyase. ^[12]^ However, both aspartate and fumaric acid are made from fossil carbon resources, such as oil and coal. Another method is the use of engineered aspartase which converts acrylic acid and ammonia to β-alanine. ^[13]^ However, acrylic acid is synthesized via the two-step oxidation method of propylene, which also relies on petrochemical resources. The third one is through natural lysine 2,3-aminomutase (EC 5.4.3.2), which was found to have a promiscuous activity on α-alanine to synthesize β-alanine ^[14]^, called aminomutation. However, this S-adenosyl methionine (SAM) ^[12b]^ dependent enzyme and its mutants have very low activities on its non-natural substrate. Furthermore, it requires expensive cofactors and strict anaerobic conditions for its proper assembly of [4Fe-4S] clusters, limiting its potential for industrial application. ^[15]^

Herein we designed an *in vitro* biotransformation pathway composed of two cascade decarboxylases, i.e., aspartate β-decarboxylase (ABD) and AAD, and implemented aminomutation from α-alanine to β-alanine by coupling the two decarboxylases. Interestingly, we also discovered a new chemical reaction of ABD that can fix CO_2_ directly. *In vitro* CO_2_ self-sufficiency and intramolecular amino-editing were achieved in this designed process with 100% carbon atom economy.

## Results

### Design *of in vitro* β-alanine biosynthesis

α-Alanine has been fermented from glucose and ammonia by an engineered *E. coli* on a large industrial scale. ^[16]^ However, lysine 2,3-aminomutase (EC 5.4.3.2) cannot convert α-alanine to β-alanine effectively. Inspired by the CO_2_ fixing reaction catalyzed by formate dehydrogenase, we hypothesized that ABD (EC 4.1.1.12) may fix CO_2_ from α-alanine to α-aspartate. The analysis of the eQuilibrator found that the equilibrium constant of the CO_2_-fixing reaction catalyzed by ABD was greater than that by formate dehydrogenase. Therefore, we designed an *in vitro* two-enzyme reaction pathway consisting of ABD and AAD to implement amino-editing from α-alanine to β-alanine featuring CO_2_ self-sufficiency (**Figure 1A**). In it, ABD may catalyze the hypothetical CO_2_-fixing reaction for the biosynthesis of α-aspartate from α-alanine and CO_2_; AAD can catalyze its decarboxylation reaction of α-aspartate for the synthesis of β-alanine with release of CO_2_. The standard Gibbs free energy change (ΔG′°) of each step is shown in **Figure 1B**. The overall ΔG′° of the two cascade reactions was calculated to be −5.1 kJ mol^−1^ at pH 7.5, suggesting β-alanine formation preferred. Evidently, the critical step in this two-step reaction is the carboxylation, so the choice of a suitable decarboxylase and optimal circumstances that favor its reverse reaction is essential.

**Figure 1.**
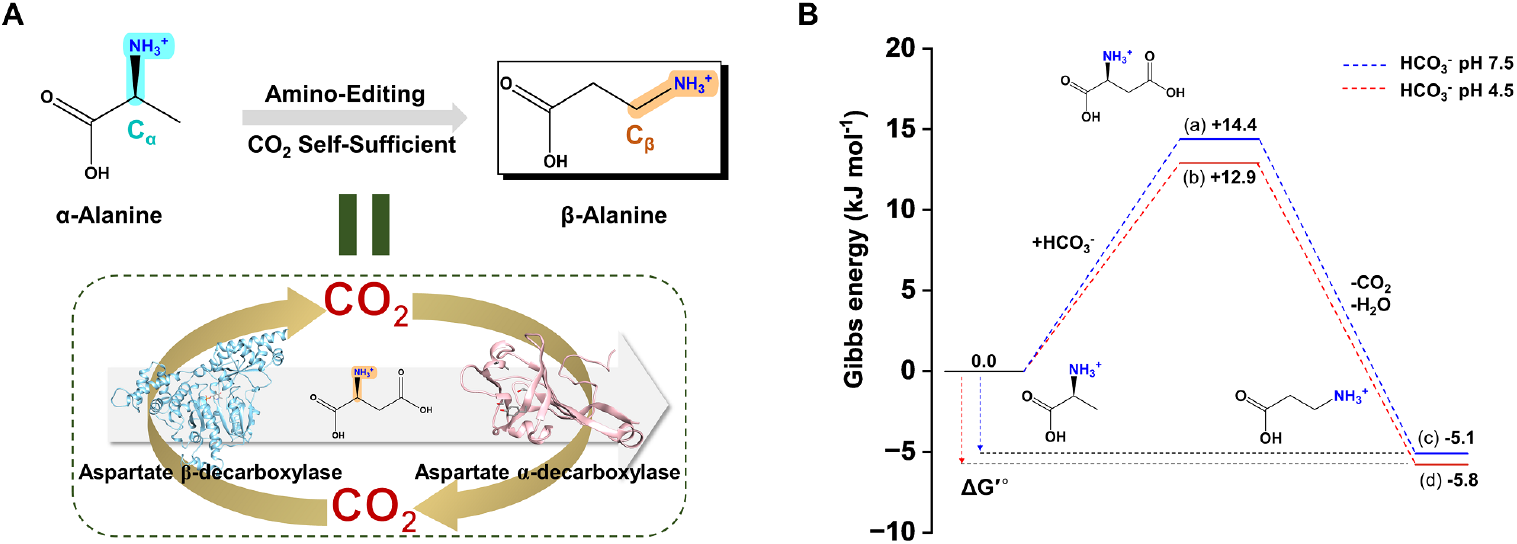
Scheme of the proposed enzymatic biosynthesis of β-alanine from α-alanine and Gibbs free energy analysis on the involved reactions. (A) The amino-editing of α-alanine by CO_2_ self-sufficient aspartate β-decarboxylase (ABD) and aspartate α-decarboxylase (AAD). (B) Standard Gibbs free energy changes for each cascade reaction under conditions of ionic strength of 0.25 M and pH 7.5 (blue) or pH 4.5 (red).

### Mining of ABD for CO_2_ fixing reaction

ABD catalyzes the β-decarboxylation of α-aspartate to α-alanine and CO_2_. This enzyme belongs to class I pyridoxal 5’-phosphate (PLP)-dependent aspartate aminotransferase superfamily, with a low-level nonspecific α-amino acid transamination activity. ^[17]^ Before this study, there was no report pertaining to the CO_2_-fixing reaction catalyzed by ABD. As described above, our hypothesis is ABD may be able to catalyze the CO_2_-fixing reaction to synthesize α-aspartate from α-alanine and CO_2_. Due to biomanufacturing advantages of thermostable enzymes ^[18]^, we screened five enzymes from thermophiles with 2,000 putative ABD-encoding sequences in the family of ABD (0A8J7M890), that is, *Thermohalobaculum xanthum* ABD (TxABD, Uniprot A0A8J7M890), *Thermococcus* sp. MV11 (TmABD, Uniprot A0A8J8A898), *Acinetobacter radioresistens* DSM 6976 (ArABD, NCBI BBL20616.1), *Clostridium thermobutyricum* (CtABD, Uniprot N9Y2J5), and *Thermoactinospora* sp. (TsABD, Uniprot A0A8J8HAW7). Multiple sequence alignment results showed their sequence similarity from 42% to 53%. The sequences near the coenzyme PLP binding lysine were highly conserved and the conserved sequence motif (signature: SXSKYFGXTGWRLG) was the aminotransferase class-I PLP attachment site (**Figure S1A**). We further compared their similarities using phylogenetic analysis. The resultant neighbor-joining tree clustered the ABDs in two independent branches with thermophilic-derived ABDs (TsABD, CtABD, and TxABD) and thermostable ones (ArABD and TmABD) (**Figure S1B**).

The five ABD genes were synthesized, expressed in *E. coli*, and the corresponding enzymes were purified (**Figure S2**). In order to optimize the *in vitro* CO_2_ fixing system, we firstly measured the specific activities of those ABDs in converting α-alanine to α-aspartate at pH 7.5 (in HEPES buffer), containing 100 mM α-alanine, 1 mM pyridoxal 5’-phosphate (PLP), 100 mM NaHCO_3_, and 1 g/L ABD. TsABD showed a higher specific activity of carboxylation (**Figure S3**). Additionally, as ΔG′° indicated that an acidic environment was favorable to the carboxylation reaction (**Figure 1B**), we also analyzed the specific activities of carboxylation at pH 4.5 (in the citric acid buffer). The activities of all ABDs were increased in the acidic environment, still with TsABD being the highest (**Figure S3**). We further confirmed the α-aspartate produced in the reaction by mass spectrometry analysis (**Figure S4**). Taking all the above together, the reversing amino acid decarboxylase reaction was validated for CO_2_-fixing reaction, and the TsABD was selected for the carboxylation of α-alanine in the following study.

### Optimization of α-alanine carboxylation

For the carboxylation reaction optimization, the effect of ions was investigated, such as, CO^2+^, Mn^2+^, Ni^2+^, Ca^2+^, Cu^2+^, Zn^2+^, Fe^3+^, Mg^2+^, Fe^2+^ and their respective concentrations (0 mM, 1 mM, and 5 mM), to the reaction system. The results showed an inhibitory effect of those ions compared with the control (**Figure S5**). Next, we measured the specific activities at a range of pH 2.5-8.0, in different buffers (citrate buffer, phosphate buffer, and HEPES buffer). The highest activity of carboxylation was obtained at pH 3.5 (in the citric acid buffer). Besides, TsABD showed a higher activity in an acidic pH environment, beyond this range, the specific activities of carboxylation decreased, which was also consistent with what the ΔG′° indicated (**Figure S6**). To optimize the temperature, we first assayed the carboxylation activity of TsABD at pH 7.5 in a range of 37°C-80°C. The carboxylation activity gradually increased with a higher temperature and reached the highest at 75°C (**Figure S7A**). However, when the activity was determined at the optimal pH 3.5 in a range of 37°C-70°C, the carboxylation activity reached the maximum at 55°C (**Figure S7B**). The results indicated that the pH value may have a significant effect on CO_2_ fixation to shift the equilibrium towards carboxylation. Therefore, the optimal conditions for the carboxylation reaction of TsABD were determined as pH 3.5 and 55°C.

Subsequently, the kinetic analysis was performed at the above optimal conditions. The reaction system included 100 mM citric acid buffer (pH 3.5), 1 mM PLP, and 1 g/L TsABD with a gradient concentration of α-alanine (0-200 mM) and NaHCO_3_ (0-200 mM). From curve fitting of the initial reaction velocities, TsABD showed almost identical *k*_*m*_ value for the α-alanine (47.6 mM) substrate and NaHCO_3_ (45.7 mM) substrate, and the same as *k*_*cat*_ value (**Figure S8**). Furthermore, we studied the multi-substrate binding mechanism with a range of α-alanine concentration gradient (0-200 mM) by fixing NaHCO_3_ at 10 mM, 50 mM, or 100 mM. The curve was plotted with Lineweaver-Burk double reciprocal graphing, the reciprocal of the reaction rate (1/v) as the ordinate and the substrate concentration (1/[s]) as the abscissa. For sequential mechanisms in bi-substrate reactions, Lineweaver-Burk plots at varying A and different fixed values of B give a series of intersecting lines. When intersecting on the X-axis, the binding mechanism is the random binding mechanism. As shown in **Figure S9**, the bi-substrate binding mechanism of TsABD was random mechanism, different from the ping-pong mechanism that PLP-dependent aminotransferases (EC 2.6.1.x) followed. ^[19]^ In random sequential reactions, the substrates and products are bound and then released in no preferred order, or “random” order. The two substrates of α-alanine and NaHCO_3_ do not affect each other and bind first when they contact with the enzyme randomly.

### Proof-of-concept of the proposed *in vitro* biosynthesis pathway

AAD, consisting of Pyruvoyl-type and PLP-type, catalyzes the α-decarboxylation of α-aspartate to β-alanine and CO_2_, which is the major route of β-alanine production in bacteria. To catalyze the biotransformation of the intermediate α-aspartate into β-alanine, we selected AADs by BRENDA enzyme database (https://brenda-enzymes.org/) search and gene mining of thermostable and thermophilic-derived enzymes. Six candidates of AAD derived from *Corynebacterium glutamicum* ATCC 13032 (CgAAD, Uniprot Q9X4N0), *Bacillus subtilis* 168 (BsAAD, Uniprot P52999), *Archaeoglobus fulgidus* (AfAAD, Uniprot A0A075WHK3), *Pyrococcus furiosus* (PfAAD, Uniprot Q8U1P6), *Thermococcus kodakarensis* (TkABD, Uniprot Q5JJ82), and *Thermus thermophilus* (AfAAD, Uniprot Q72L22) were selected. We investigated these similarities using the phylogenetic analysis based on DNA sequences. The resultant neighbor-joining tree clustered the AADs in two independent branches, i.e., Pyruvoyl-type AAD (BsAAD, TtAAD, and CgAAD) and PLP-type AAD (AfAAD, PfAAD, and TkAAD) (**Figure S10**). Next, the two types of AAD genes were amplified, expressed in *E. coli*, and the corresponding enzymes were purified (**Figure S11**). The AAD activities were determined at pH 7.5 (in the HEPES buffer). CgAAD showed the highest decarboxylation activity (**Figure S12**). Thus, coupled with TsABD, the new cascade biotransformation of β-alanine was achieved (**Figure 2A**).

**Figure 2.**
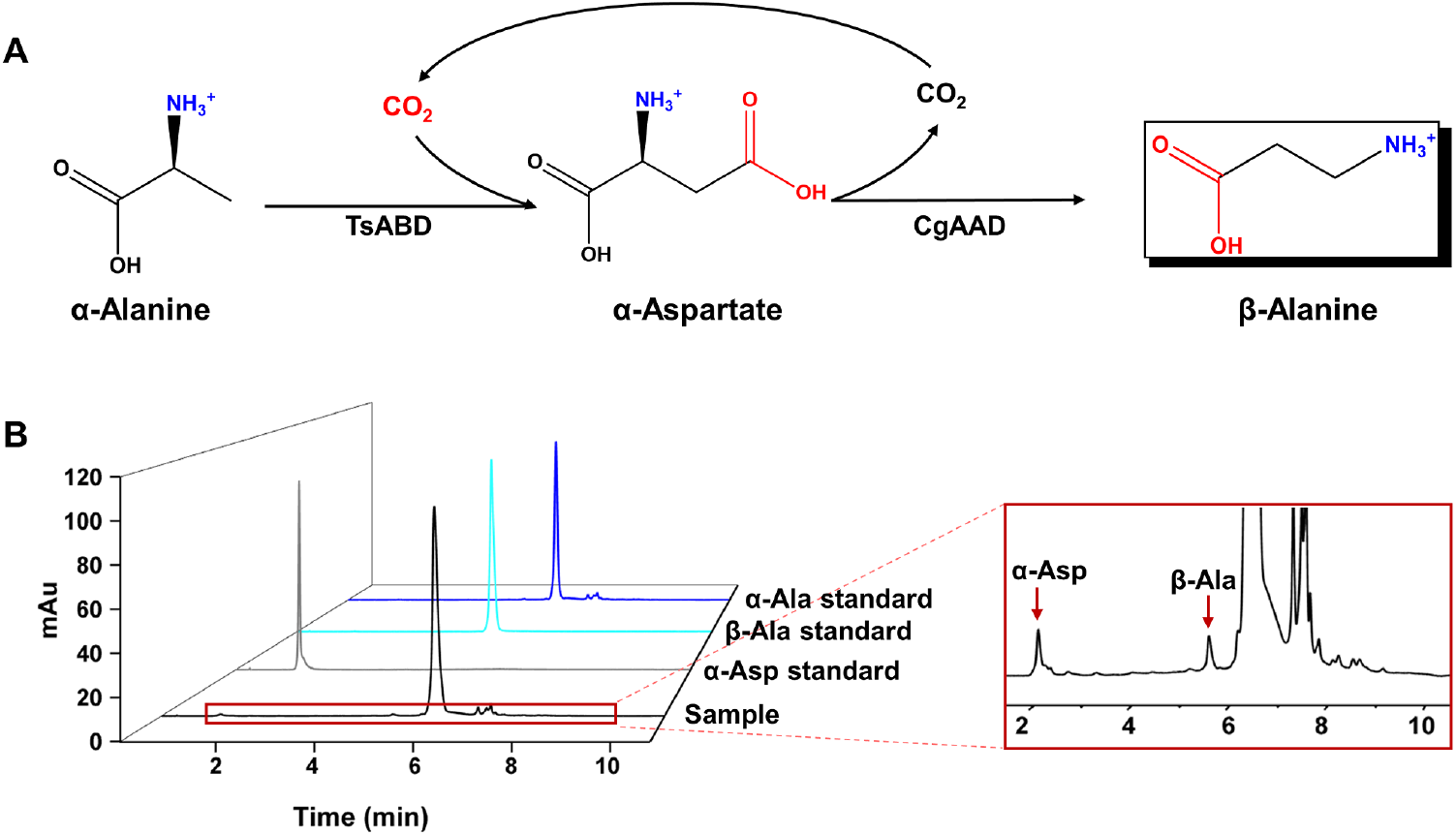
Validation of the new β-alanine biotransformation pathway. (A) Scheme of the bioctransformation of β-alanine from α-alanine via α-aspartate. The α-aspartate served as the intermediate of α-alanine carboxylation by TsABD (ABD from *Thermoactinospora* sp.) and was subsequently converted using CgAAD (AAD from *Corynebacterium glutamicum* ATCC 13032). (B) *In vitro* cascade synthesis of β-alanine at 37°C for 24 h in 100-mM HEPES buffer (pH 7.5), containing 100 mM α-alanine, 1 mM PLP, 100 mM NaHCO_3_, 1 g/L TsABD, and 1 g/L CgAAD. α-Alanine, α-aspartate, and β-alanine in the reaction were detected using HPLC.

To validate the new *in vitro* biotransformation pathway, based on the above results, both TsABD and CgAAD were employed for the biosynthesis of β-alanine from α-alanine with a co-substrate NaHCO_3_. We initially combined the enzymes in a weight ratio of 1:1 (mg: mg) and reacted at 37°C with 100 mM HEPES buffer (pH 7.5), 100 mM α-alanine, 1 mM PLP, 100 mM NaHCO_3_, 1 g/L CgAAD, and 1 g/L TsABD. After 24 h, the substrate (α-alanine), intermediate (α-aspartate) and product (β-alanine) in the reaction mixture were all determined by HPLC (**Figure 2B**) and further confirmed by GC-MS (**Figure S13**). Both methods were able to detect the intermediate α-aspartate and the target product β-alanine. Therefore, the hypothetical *in vitro* biotransformation pathway from α-alanine to β-alanine was validated for the first time. This pathway by coupling the two decarboxylases implemented intramolecular amino editing from α-alanine to β-alanine with 100% carbon atom economy.

### Optimization of the cascade conditions for β-alanine production

In order to optimize the novel *in vitro* system for β-alanine production, we next determined the pH and temperature conditions of the cascade reaction. The optimum pH for the biocatalytic synthesis was measured at 37°C with 100 mM α-alanine, 1 mM PLP, 100 mM NaHCO_3_, 1 g/L CgAAD, and 1 g/L TsABD in 100 mM citrate buffer (pH 2.5-7.0). The optimum temperature for the biosynthesis of β-alanine was determined at pH 4.5 (100 mM citrate buffer) with 100 mM α-alanine, 1 mM PLP, 100 mM NaHCO_3_, 1 g/L CgAAD, and 1 g/L TsABD in a 37°C-70°C range. In contrast to the results above that AAD was used to decarboxylate α-aspartate at alkaline conditions, the results here showed the highest β-alanine biocatalytic activity at pH 4.5 and 55°C (**Figure 3A and 3B**), suggesting a preference for the rate-limiting carboxylation reaction in this two-step cascade reaction. For further optimization, a series of enzyme ratios of TsABD/CgABD were designed. After reaction for 2 h, the highest β-alanine production was obtained with a TsABD/CgABD weight ratio of 10:0.1 (**Figure 3C**). The results indicated that an elevated ABD concentration resulted in a significantly enhanced level of the final β-alanine, demonstrating the rate-limiting role of the decarboxylase for the coupled reaction. Therefore, the optimal conditions were at pH 4.5, 55°C, and an enzyme ratio of 10:0.1 (TsABD/CgABD) for the β-alanine production. A higher amount of ABD ensured that there was sufficient enzyme at the end of the reaction to balance the rate-limiting step. The final β-alanine biosynthesis reaction system comprised 100 mM citric acid buffer (pH 4.5), 100 mM α-alanine, 1 mM PLP, 100 mM NaHCO_3_, 0.1 g/L CgAAD, and 10 g/L TsABD. The α-alanine, α-aspartate, and β-alanine in the reaction mixture were confirmed by HPLC (**Figure S14**). However, with the extension of time, the intermediate α-aspartate reached its highest at 24 h and was gradually converted to β-alanine (**Figure 3D**), indicating that deactivation of at least one of the enzymes may limit the biocatalytic reaction with longer incubation time. The highest β-alanine production was obtained at the optimal condition, yielding a β-alanine concentration of approximately 0.98 mM at 120 h. In this study, the lumped specific activity of the two enzymes from α-alanine to β-alanine is 34.72 nmol/min/mg enzyme, were 165- and 2.89-fold higher than those catalyzed by LAM without and with ethylamine as an activator ^[14c]^.

**Figure 3.**
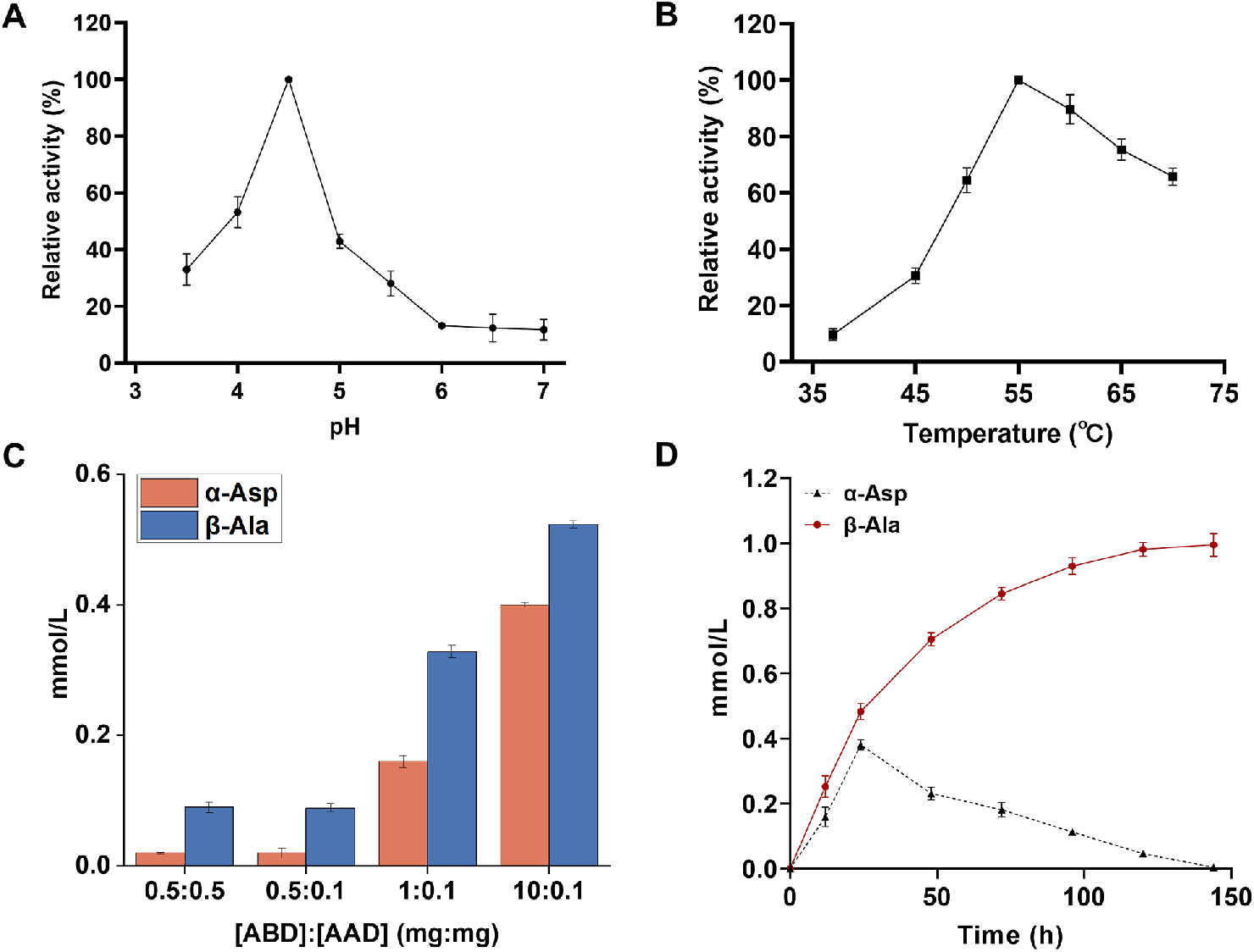
Optimization of enzymatic reaction conditions for β-alanine production. Optimization of pH (A), temperature (B), and the ratio of enzymes (C) in the *in vitro* cascade reaction. (D) *In vitro* cascade synthesis of β-alanine at optimal reaction conditions (at 55°C in 100 mM citric acid buffer (pH 4.5), containing 100 mM α-alanine, 1 mM PLP, 100 mM NaHCO_3_, 10 g/L TsABD, and 0.1 g/L CgAAD). The α-alanine, α-aspartate, and β-alanine in the reaction was detected by HPLC. Error bars indicated s.d. from triplicate measurements.

These data suggested that this two-enzyme system composed of two decarboxylases was superior to one-enzyme LAM in terms of specific activity.

Further, to ensure that CO_2_ was recycled in the biosynthesis of β-alanine, ^13^C-NaHCO_3_ labeling approach was applied to verify CO_2_ self-sufficiency. To this end, the cascade reaction samples incubated with 100 mM ^13^C-NaHCO_3_ at 96 h were collected and analyzed using GC/Q-TOF-MS (**Figure S15**). After assimilated ^13^C-NaHCO_3_ was converted to the intermediate α-aspartate, the ^13^C-labeling α-aspartate lost a molecule of CO_2_ that was not labeled and ^13^C-labeling β-alanine was synthesized. Subsequently, the CO_2_ (not labeled) entered into the cascade reaction, resulting in that both ^13^C-labeling β-alanine and not labeled β-alanine were both detected in the reaction mixture as shown in **Figure S15**. Accordingly, the intramolecular amino editing from α-alanine to β-alanine with CO_2_ self-sufficiency by cascade two decarboxylases was demonstrated.

Molecular editing of highly functionalized compounds, which are attached to functional groups of alcohols, amines, or carboxylic acids, refers to the process of modifying molecules at the atomic level by inserting, deleting, or exchanging atoms. ^[20]^ It is a powerful tool for modifying the properties of a molecule to improve its functionality, effectiveness, or safety by creating another (new) compound. To date, molecular editing has a broad range of applications in many fields such as drug discovery, natural product synthesis, enzyme engineering, materials science, and others. ^[21]^ Moreover, aminomutation is also a crucial area of molecular editing for exchanging a amino group. For example, aminomutase can catalyze the intramolecular rearrangement of amino acids, converting α-amino acids to their corresponding β-amino acids. ^[15a]^ Enhancing the activity of alanine 2,3-aminomutase for industrial β-alanine production from α-alanine could be economically attractive.

In this work, we developed a novel biochemical pathway for the intramolecular editing of β-amino acids from α-amino acids. By reconstructing biotransformation pathway of β-alanine from α-alanine, a green and efficient molecular editing method referred to *in vitro* CO_2_ self-sufficiency with 100% carbon atom economy was achieved. Importantly, we have presented the first amino acid decarboxylase for the CO_2_ fixation. In fact, using two enzymes to perform a single enzyme function may increase the efficiency of the reaction, provide more flexibility in the reaction conditions and offer better specificity for the reaction. For example, trehalose synthase (TreS) for trehalose production from maltose, and its biosynthesis can be replaced by the trehalose-6-phosphate synthase (TPS)/trehalose-6-phosphate phosphatase (TPP) pathway. The latter consists of a two-step catalysis by the TPS and TPP via UDP-glucose and glucose 6-phosphate, which is more efficient and most widely used in industry to date. ^[22]^ In addition, xylose reductase and xylitol dehydrogenase with NADH self-sufficiency was validated for the conversion of xylose to xylulose in yeast, which was catalyzed by xylose isomerase in bacteria. ^[23]^ These examples illustrate how organisms can use different enzymes and pathways to accomplish the same overall reaction.

## Conclusion

This work demonstrated the proof-of-concept regarding a novel route for the intramolecular editing of β-amino acids from α-amino acids. By reconstructing biotransformation of β-alanine from α-alanine, we discovered the first amino acid decarboxylase capable of fixing CO_2_ directly. Despite the presumed irreversible nature of decarboxylation, coupling of the reverse enzymatic reaction with a decarboxylase was sufficient to shift the chemical equilibrium, leading to thermodynamically favorable production of β-alanine. In addition, *In vitro* CO_2_ self-sufficiency and 100% carbon atom economy were achieved in the novel catalysis pathway. Taken together, our findings represent a significant advancement in the field of biosynthesis and sustainable chemistry, with important implications for the industrial production of β-alanine and the field of intramolecular editing of other β-amino acids. The discovery of a CO_2_-fixing amino acid decarboxylase could open up new possibilities for the development of sustainable and energy-efficient chemical processes, as well as for the creation of novel amino acid derivatives with potential biological activities.

## Supporting information

Supplemental Experimental Procedures, Figure S1-S15 and Table S1-S5.

## Acknowledgments

This work was supported by the Major Project of Haihe Laboratory of Synthetic Biology (22HHSWSS00015), the National Key Research and Development Program of China (2022YFA0912300), and Tianjin Synthetic Biotechnology Innovation Capacity Improvement Project, China (TSBICIP-CXRC-067).

## Entry for the Table of Contents

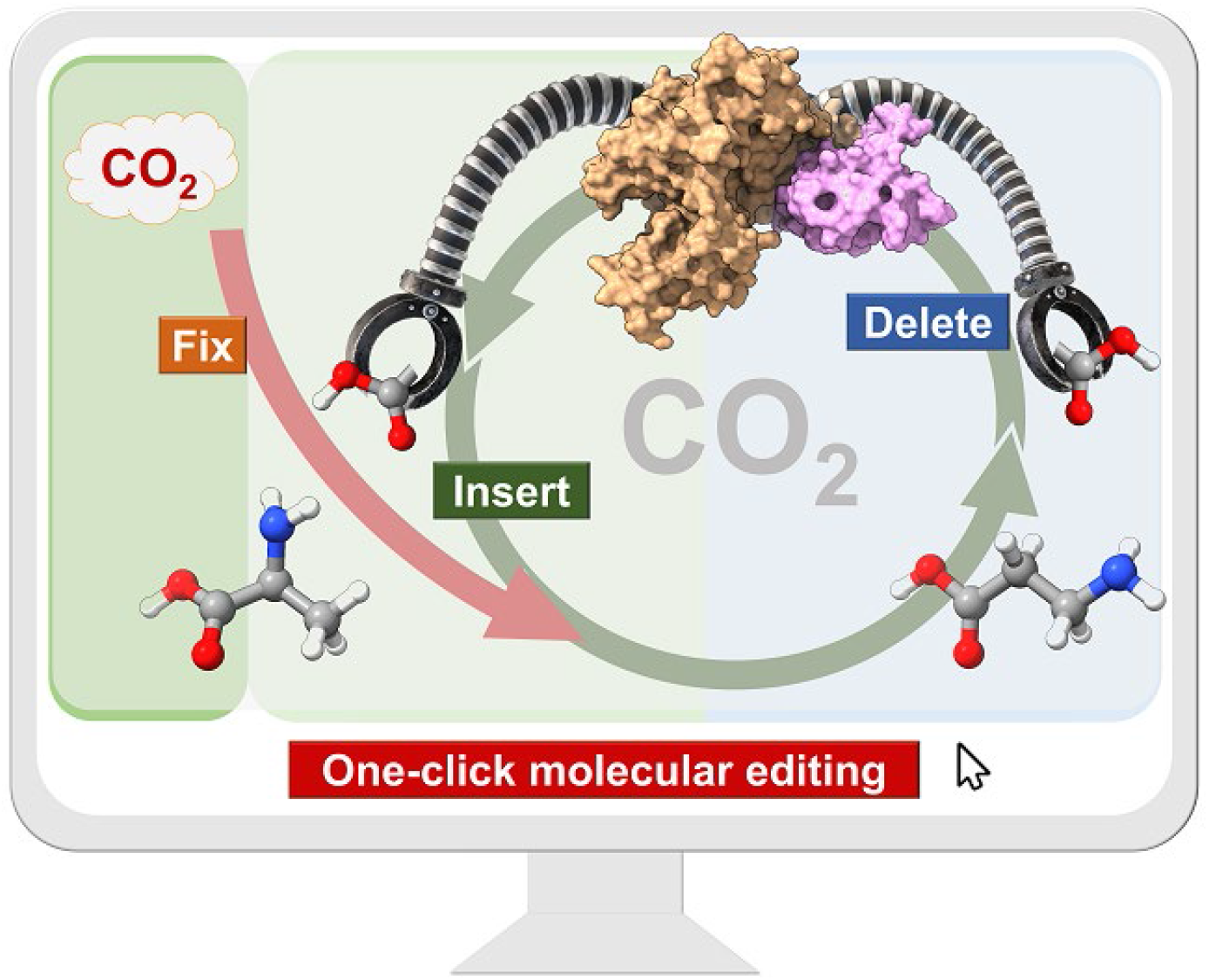

A novel one-click intermolecular aminomutation from α-position of α-amino acids to β-position of β-amino acids was implemented by two cascade decarboxylases. The first amino acid decarboxylase which can fix CO_2_ directly. Furthermore, the new biotransformation pathway features *in vitro* CO_2_ self-sufficiency and 100% carbon atom economy.

## Notes

### Competing Interest Statement

The authors have declared no competing interest.

