## Supplemental Experimental Procedures, Figure S1-S15 and Table S1-S5. for "Aminomutation catalyzed by CO_2_ self-sufficient cascade amino acid decarboxylases"

#### Materials and chemicals

All chemicals were reagent grade or higher, purchased from Sigma-Aldrich (St. Louis, MO, USA), Sinopharm (Shanghai, China), or Aladdin (Shanghai, China), unless otherwise specified. PrimeSTAR Max DNA Polymerase from Takara (Tokyo, Japan) was used for the PCR reactions.

#### Plasmids and strains

After sequence alignment and gene mining, five different sources of aspartate β-decarboxylase (EC 4.1.1.12, ABD) and six different sources of aspartate α-decarboxylase (EC 4.1.1.11, AAD) were selected. The coding sequences for ABDs from *Thermohalobaculum xanthum* ABD (TxABD, Uniprot A0A8J7M890), *Thermococcus* sp. MV11 ABD (TmABD, Uniprot A0A8J8A898), *Acinetobacter radioresistens* DSM 6976 (ArABD, NCBI BBL20616.1), *Clostridium thermobutyricum* (CtABD, Uniprot N9Y2J5), *Thermoactinospora* sp (TsABD, Uniprot A0A8J8HAW7) were synthesized by GENEWZ (Suzhou, China) with codon optimization and cloned into the vector pET28a (+) between the *Nde* I and *Eco*R I restriction sites.

The AADs from *Corynebacterium glutamicum* ATCC 13032 (CgAAD, Uniprot Q9X4N0), *Bacillus subtilis* 168 (BsAAD, Uniprot P52999), *Archaeoglobus fulgidus* (AfAAD, Uniprot A0A075WHK3), *Pyrococcus furiosus* (PfAAD, Uniprot Q8U1P6), *Thermococcus kodakarensis* (TkABD, Uniprot Q5JJ82), *Thermus thermophilus* (TtAAD, Uniprot Q72L22) were all genomically cloned and assembled between the *Nde* I and *Xho* I restriction cleavage sites of the vector pET28a (+). For example, the insertion fragment containing the *CgAAD* gene was amplified from the *Corynebacterium glutamicum* ATCC 13032 genome using a primer pair of *CgAAD*-IF and *CgAAD*-IR, while the pET28a vector backbone was amplified with a primer pair of *CgAAD*-VF and *CgAAD*-VR (Table S1). The amplified insertion fragment was then cloned into the vector via prolonged overlap extension PCR (POE-PCR). The primers and plasmids used in this study are listed in Table S1 and Table S2. The host strain *E. coli* TOP10 was used for DNA manipulation. The host strain *E. coli* BL21 (DE3) was used for recombinant protein expression. The strains used in this study are listed in Table S3.

#### Protein expression and purification

Bacteria were grown with shaking in lysogeny broth medium supplemented with 50 μg/mL kanamycin at 37 °C. When the OD600 reached 0.6-0.8, 0.1 mM isopropyl-β-D-thiogalactopyranoside (IPTG) was added to induce the overexpression of enzymes at 16 °C for 16-18 h. The bacteria were then collected by centrifugation at 6,000 rpm for 15 min at 4 °C and resuspended in 50 mM HEPES buffer (pH 7.4) containing 0.1 M NaCl. After ultrasonic disruption, the supernatant was collected by centrifugation at 8000 rpm for 30 min at 4 °C and the cell pellets were resuspended in the same buffer to a final OD600 of 50, the His6-tagged enzymes were trapped on Ni-NTA superflow resin (Qiagen, Hilden, Germany). The purity of the recombinant proteins was examined by SDS-PAGE, and the protein concentration was determined using the Bradford method with bovine serum albumin.

**HPLC analysis**

The levels of α-alanine, α-aspartate, and β-alanine were determined using high-performance liquid chromatography (HPLC) with an AJS-02 amino acid-specific analytical column-C18 column (4.6 mm×150 mm, 3 μm, Shimadzu) and UV-detector. Prior to subjecting the samples to the column, they were derivatized using boric acid and *o*-phthalaldehyde (OPA).

The gradient elution from buffer A (10 mM $Na_2HPO_4$•$12H_2O$ and 10 mM $Na_2B_4O_7$•$10H_2O$, pH 8.2) to buffer B (acetonitrile, methanol, and water with a volume ratio of 45: 45: 10) was performed at 50°C and 1.6 mL $min^{-1}$ flow rate. The retention times for α-aspartate, β-alanine, and α-alanine were 1.5 min, 5.8 min, and 6.9 min, respectively.

**Enzyme activity assay**

The activity of aspartate β-decarboxylase (ABD) on α-alanine was assayed at 37°C for 2 h. The reaction system (1 mL) consisted of 100 mM HEPES buffer (pH 7.5) or 100 mM citric acid buffer (pH 4.5), 100 mM α-alanine, 1 mM pyridoxal 5'-phosphate (PLP), 100 mM $NaHCO_3$, and 1 g/L ABD. The reaction was stopped by adding 0.1 mL of 1 M NaOH. One unit of ABD activity was defined as the amount of enzyme producing 1 μmol of α-aspartate per minute.

The activity of aspartate α-decarboxylase (AAD) on α-aspartate was assayed at 37°C for 5 min. The reaction system for pyruvoyl-type AAD (1 mL) comprised 100 mM HEPES buffer (pH 7.5), 100 mM α-aspartate, and 0.1 g/L AAD. The reaction system for PLP-type AAD (1 mL) consisted of 100 mM HEPES buffer (pH 7.5), 100 mM α-aspartate, 1 mM PLP, and 0.1 g/L AAD. The reaction was stopped by adding 0.1 mL of 1 M NaOH. One unit of AAD activity was defined as the amount of enzyme producing 1 μmol of β-alanine per minute.

**Enzymatic property assay**

The optimum ion for ABD was determined at 37°C for 2 h. The reaction system (1 mL) comprised 100 mM HEPES buffer (pH 7.5), 100 mM α-alanine, 1 mM PLP, 100 mM $NaHCO_3$, and 1 g/L TsABD with 0 mM iron (control) or 1 mM iron ($CO^{2+}$, $Mn^{2+}$, $Ni^{2+}$, $Ca^{2+}$, $Cu^{2+}$, $Zn^{2+}$, $Fe^{3+}$, $Mg^{2+}$, $Fe^{2+}$) or 5 mM iron ($Fe^{3+}$, $Mg^{2+}$, $Fe^{2+}$).

The optimum pH for ABD was determined at 37°C for 2 h. The reaction system (1 mL) comprised 100 mM α-alanine, 1 mM PLP, 100 mM $NaHCO_3$, and 1 g/L TsABD in different buffers, including 100 mM citrate buffer (pH 2.5-7.0), 100 mM phosphate buffer (pH 6.0–8.0), and 100 mM HEPES buffer (pH 7.0–

8.0).

The optimum temperature for ABD was determined at pH 7.5 (100 mM HEPES buffer) or pH 3.5 (100 mM citrate buffer) for 2 h. The reaction system (1 mL) consisted of 100 mM α-alanine, 1 mM PLP, 100 mM $NaHCO_3$, and 1 g/L TsABD in a range of 37-80°C (pH 7.5) or 37-70°C range (pH 3.5).

The reaction mixture was stopped by adding 0.1 mL of 1 M NaOH. The samples were subjected to an HPLC analysis. The curves were fitted using the Origin 8.0 software.

**Enzyme kinetic assay**

The Michaelis-Menten kinetics of ABD on α-alanine and $NaHCO_3$ were assayed at 55°C for 2 h (optimal reaction conditions). The reaction system (1 mL) comprised 100 mM citric acid buffer (pH 3.5), 1 mM PLP, and 1 g/L TsABD with varying concentrations of α-alanine (0-200 mM) and $NaHCO_3$ (0-200 mM). The reaction mixture was stopped by adding 0.1 mL of 1 M NaOH. The $k_m$ and $k_{cat}$ values were calculated by fitting the curves to the Michaelis-Menten equation using Origin 8.0 software.

**Substrate binding mechanism analysis**

The substrate binding behavior of ABD was analyzed at 55°C for 2 h. The reaction system (1 mL) consisted of $NaHCO_3$ (10 mM, 50 mM, or 100 mM), 100 mM citric acid buffer (pH 3.5), 1 mM PLP, and 1 g/L ABD with varying concentrations of α-alanine (0-200 mM). The reaction mixture was stopped by adding 0.1 mL of 1 M NaOH. The curve was plotted with Lineweaver-Burk double reciprocal graphing using Origin 8.0 software, the reciprocal of the reaction rate (1/v) as the ordinate and the substrate concentration (1/[s]) as the abscissa.

***In vitro* cascade synthesis of β-alanine**

The biocatalytic synthesis of β-alanine from α-alanine via α-aspartate was assayed at 37°C for 24 h. The reaction system (1 mL) comprised 100 mM HEPES buffer (pH 7.5), 100 mM α-alanine, 1 mM PLP, 100 mM $NaHCO_3$, 1 g/L CgAAD, and 1 g/L TsABD. The reaction was stopped by adding 0.1 mL of 1 M NaOH. The reaction mixture was detected using HPLC and GC-MS.

**GS-MS analysis of the reaction mixture**

The reaction mixture was centrifuged at 12,000 rpm for 5 min and the remaining supernatants were transferred into new centrifuge tubes and subjected to lyophilization. The dried samples were occasionally dissolved in 150 µL methoxyl amine hydrochloride (20 mg/mL in pyridine) for derivatization at 30 °C for 1.5 h. Then, 100 µL of N-methyl-N-trimethylsilyl trifluoro acetamide was added to the samples, which were then incubated at 37 °C for another 1 h. Finally, the samples were centrifuged (12,000 rpm, 10 min) and the supernatant was transferred to new GC vials.

GC-MS analysis was performed on a Thermo Scientific Orbitrap Exploris GC 240 Mass Spectrometer (Thermo Fisher, Germany). A sample volume of 1 μL was injected into an S/SL injector and separated using a TRACE 1310 gas chromatograph equipped with a TraceGOLD TG-5SILMS column (30 m × 0.25 mm, 0.25 μm film thickness). The oven temperature was programmed as follows: 60 °C for 1 min, 8 °C/min to 132 °C, 2 °C/min to 150 °C, 5 °C/ min to 185 °C, 10 °C/min to 325 °C, with a 5 min hold. Mass spectra of amino acids were collected within the mass range of 10–550 m/z at an acquisition rate of 5 spectra/s. The ion source and transfer line temperatures were set at 250 °C and 290 °C, respectively, and electron ionization was conducted at 70 eV. Agilent Mass Hunter Qualitative Analysis Software was utilized for peak detection and mass spectral deconvolution. Annotation of amino acids was carried out by comparing their mass fragmentation patterns with those in the National Institute of Standards and Technology mass spectral library (match factor >80%).

### Optimization of cascade synthesis conditions for β-alanine

The optimum pH for the biocatalytic synthesis of β-alanine from α-alanine was determined at 37°C for 2 h. The reaction system (1 mL) comprised 100 mM α-alanine, 1 mM PLP, 100 mM $NaHCO_3$, 1 g/L CgAAD, and 1 g/L TsABD in 100 mM citrate buffer (pH 2.5-7.0).

The optimum temperature for the biocatalytic synthesis of β-alanine from α-alanine was determined at pH 4.5 (100 mM citrate buffer) for 2 h. The reaction system (1 mL) comprised 100 mM α-alanine, 1 mM PLP, 100 mM $NaHCO_3$, 1 g/L CgAAD, and 1 g/L TsABD in a 37-70°C range.

The optimum enzyme ratio for the biocatalytic synthesis of β-alanine from α-alanine was determined at 55°C for 2 h. The reaction system (1 mL) comprised 100 mM citrate buffer (pH 4.5), 100 mM α-alanine, 1 mM PLP, and 100 mM $NaHCO_3$ with a different ratio of CgAAD and TsABD (0.5:0.5, 0.5:0.1, 1:0.1, and 10:0.1, mg： mg).

The reaction mixture was stopped by adding 0.1 mL of 1 M NaOH. The samples were subjected to HPLC analysis. The curves were fitted using the Origin 8.0 software.

### Production of β-alanine from α-alanine

The production of β-alanine from α-alanine via α-aspartate was assayed at optimal reaction conditions. The reaction system comprised 100 mM citric acid buffer (pH 4.5), 100 mM α-alanine, 1 mM PLP, 100 mM $NaHCO_3$, 0.1 g/L CgAAD, and 10 g/L TsABD, and was incubated at 55°C. During the reaction, samples were taken and the reaction was stopped by adding 0.1 mL of 1 M NaOH. The supernatants of samples were subjected to HPLC analysis after centrifugation (12,000 rpm, 5 min).

### Determination of $^{13}C$-labeled β-alanine

To replace the non-isotopic chemicals, 100 mM $^{13}C$-$NaHCO_3$ was added to the reaction system. The reaction was stopped by adding 1 M NaOH. The reaction mixture was centrifuged at 12,000 rpm for 5 min and the remaining supernatants were transferred into new centrifuge tubes and subjected to lyophilization. The dried samples were treated as described in the GS-MS analysis process and were finally analyzed by GC-MS.

### Figure Section

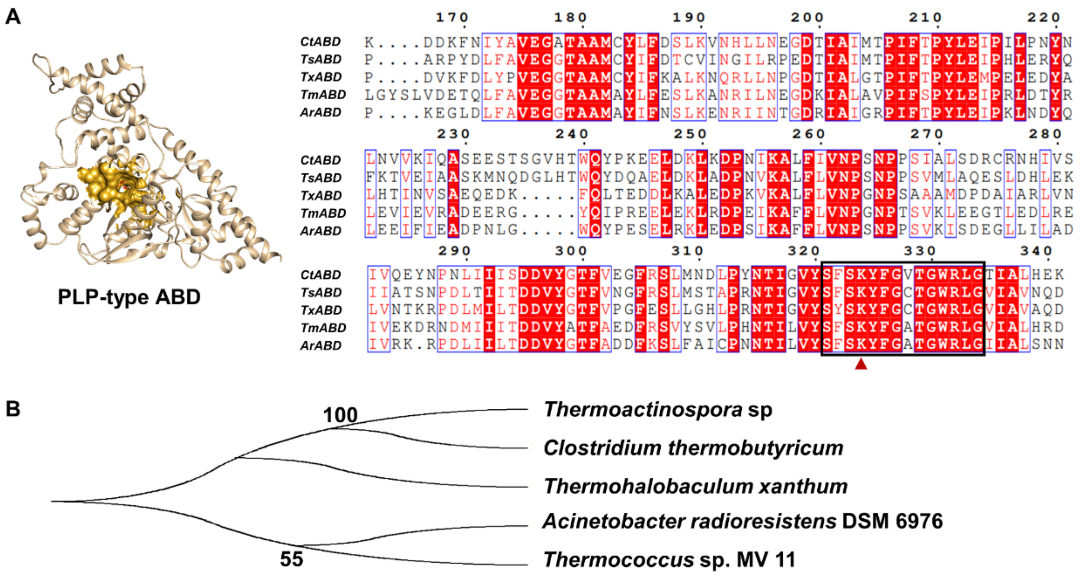

**Figure S1.** Mining of aspartate β-decarboxylase (ABD). (A) Five genes encoding aspartate β-decarboxylase were identified by database mining. The structure of CtABD was shown as cartoon (AlphaFold: AF-N9Y2J5-F1). The binding pocket of cofactor PLP was shown as surface. The boxed conserved region is the cofactor PLP attachment site. The triangular mark represents the active site Lys that binds PLP covalently during catalysis. (B) The evolutionary tree of five ABDs.

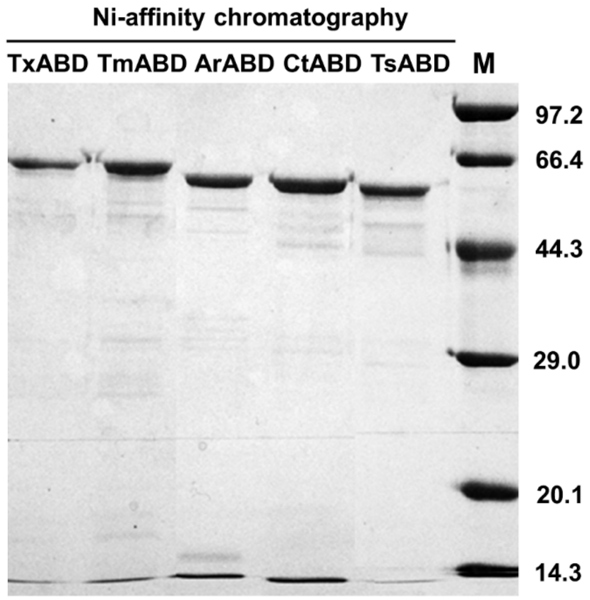

**Figure S2.** The SDS-PAGE analysis of ABDs.

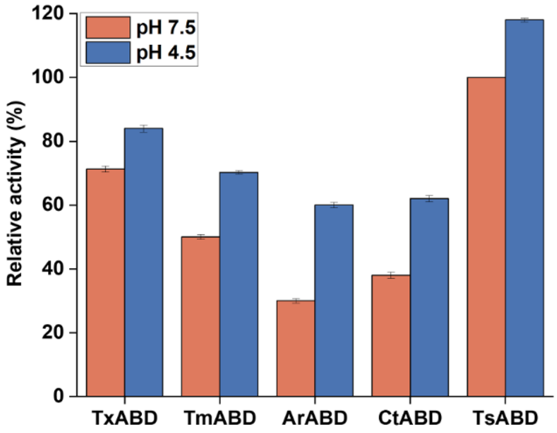

**Figure S3.** The carboxylation of α-alanine by ABDs. The relative activity of the five ABDs at pH 7.5 (in HEPES buffer) and pH 4.5 (in citric acid buffer) is shown. Error bars indicate s.d. from triplicate measurements.

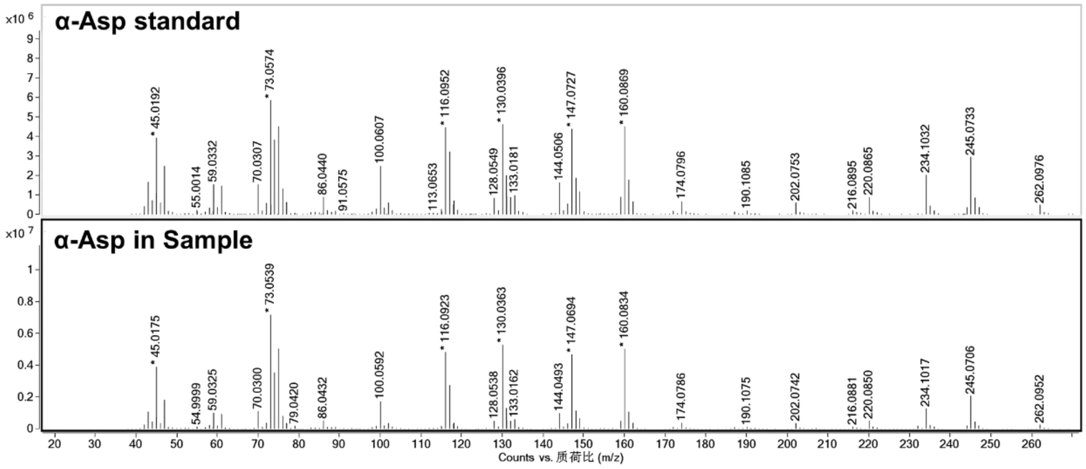

**Figure S4.** Mass spectra analysis of the synthesized α-aspartate. The activity of α-alanine carboxylation by ABD was assayed at pH 4.5 (in citric acid buffer). The reaction mixture was analyzed by GC-MS.

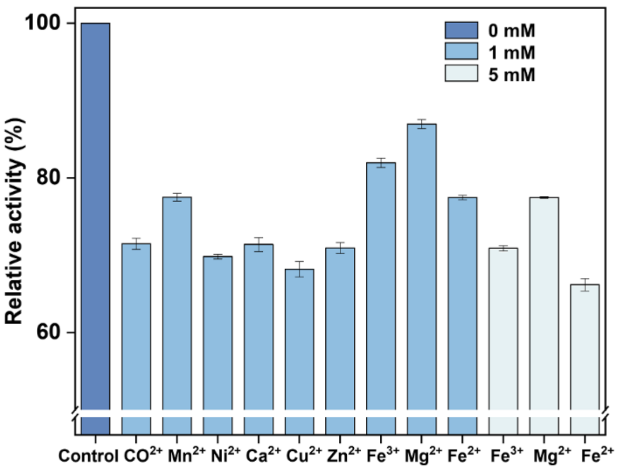

**Figure S5.** Optimization of ions in the carboxylation of α-alanine. The relative activity of TsABD at different ion conditions was shown. Error bars indicate s.d. from triplicate measurements.

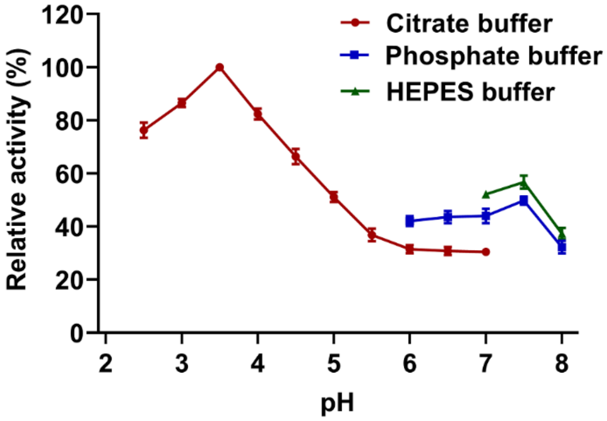

**Figure S6.** Optimization of pH in the carboxylation of α-alanine. The relative activity of TsABD at different buffer and pH conditions was shown. Error bars indicate s.d. from triplicate measurements.

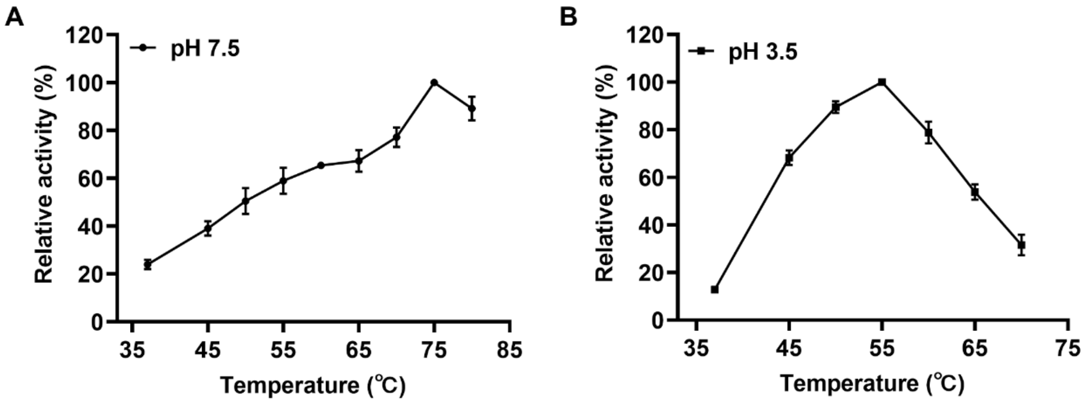

**Figure S7.** Optimization of temperature in the carboxylation of α-alanine. The relative activities of TsABD at pH 7.5 (A) and pH 3.5 (B) at different temperatures were shown. Error bars indicate s.d. from triplicate measurements.

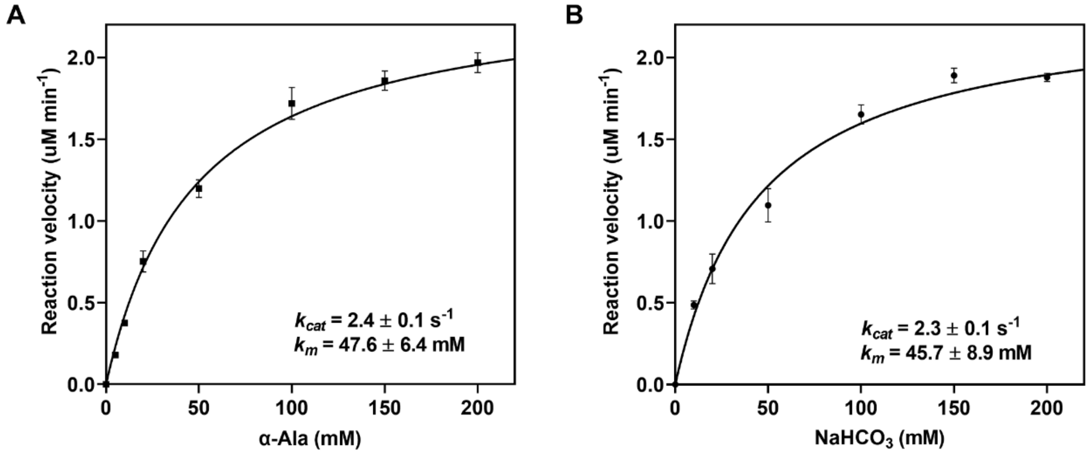

**Figure S8.** Enzymatic parameters for the decarboxylation of α-alanine. The Michaelis-Menten kinetics of the decarboxylation of α-alanine catalyzed by TsABD at optimal reaction conditions (reacted at 55°C for 2 h in 100 mM citric acid buffer (pH 3.5), contained 1 mM PLP and 1 g/L TsABD) with different concentrations of α-alanine (A) and $NaHCO_3$ (B). The $k_{cat}$ and $k_m$ are shown in the lower-right corner of each panel. Error bars indicate s.d. from triplicate measurements.

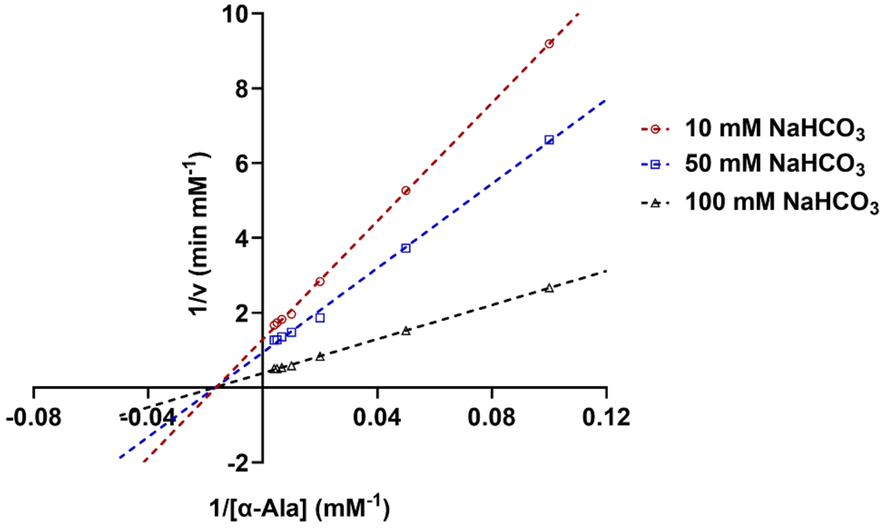

**Figure S9.** Analysis of substrate binding behaviors of ABD for the decarboxylation of α-alanine. Three groups of $NaHCO_3$ concentrations were fixed and reacted with varying concentrations of α-alanine at optimal reaction conditions (at 55°C for 2 h in 100 mM citric acid buffer (pH 3.5), containing 1 mM PLP and 1 g/L TsABD).

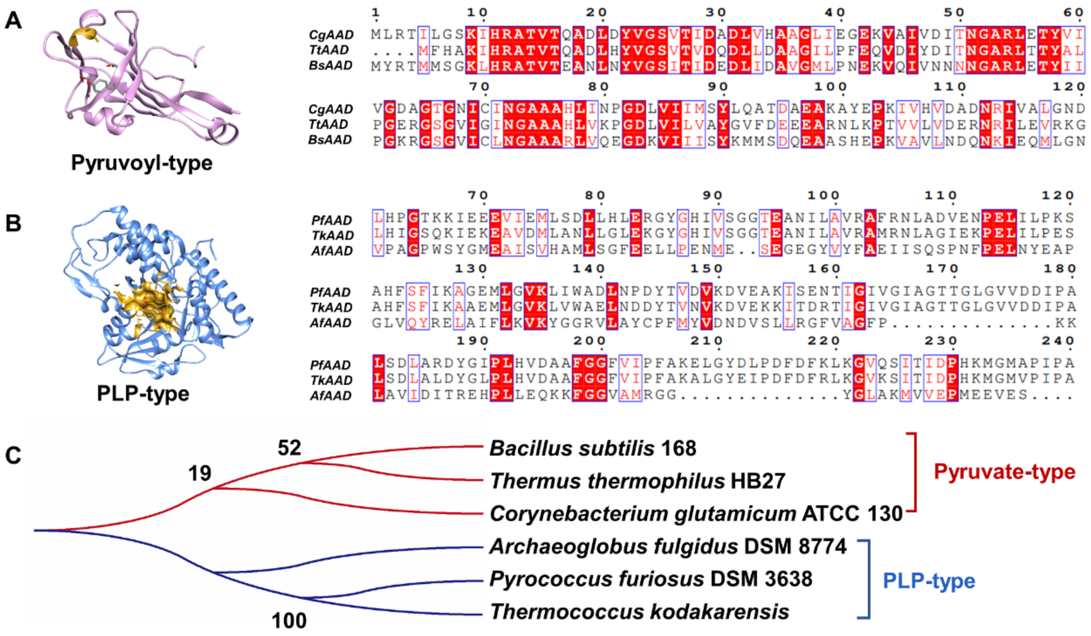

**Figure S10.** Mining of aspartate α-decarboxylase (AAD). (A) Six genes of aspartate α-decarboxylase were identified by database mining. The structure of pyruvoyl-type CgAAD (AlphaFold: AF-Q9X4N0-F1) and PLP-type PfAAD (AlphaFold: AF-Q8U1P6-F1) were shown as cartoon. The binding pocket of cofactor PLP was shown as surface. (B) The evolutionary tree of six AADs.

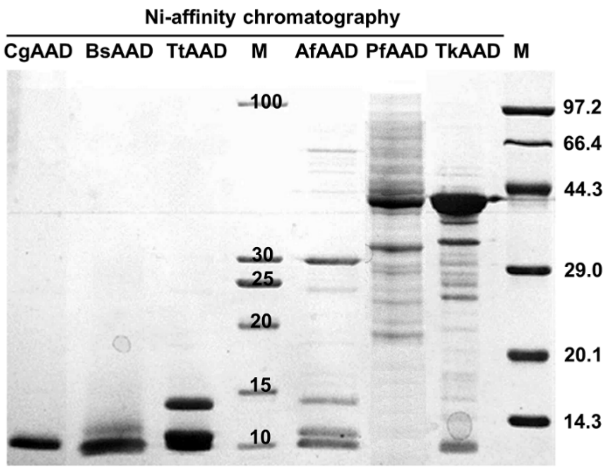

**Figure S11.** The SDS-PAGE analysis of AADs.

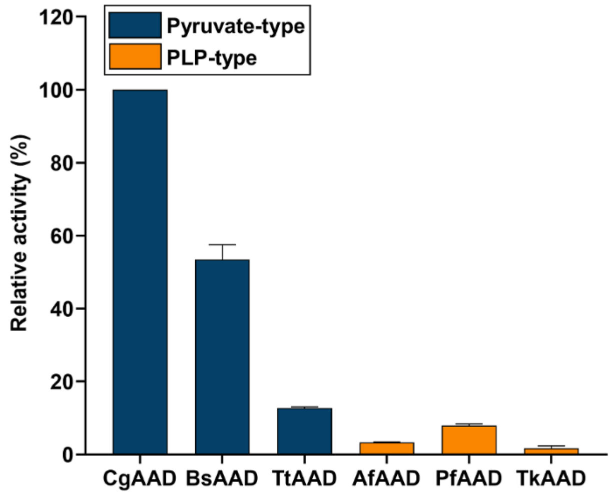

**Figure S12.** The decarboxylation of α-aspartate by AADs. The relative activity of pyruvoyl-type and PLP-type AADs at pH 7.5 (in HEPES buffer)**.** Error bars indicate s.d. from triplicate measurements.

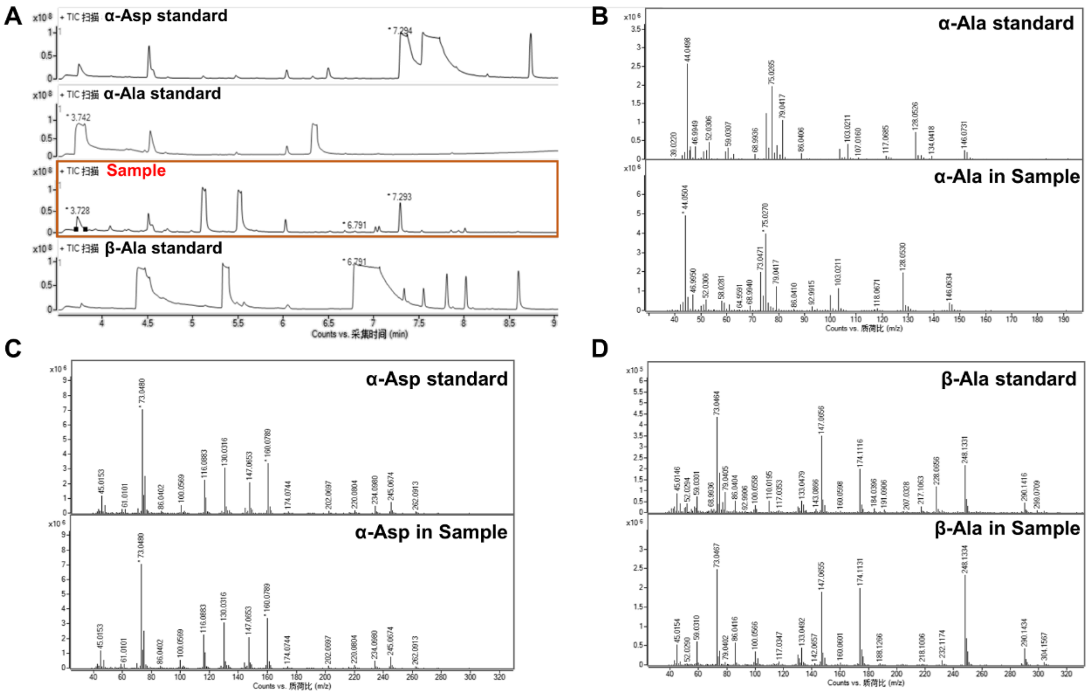

**Figure S13.** Verification of the new β-alanine biosynthesis pathway by GS-MS. The GC analysis (A) and MS analysis (B, C, and D) of α-Asp standard, α-Ala standard, β-Ala standard, and those in samples were shown. The reaction conditions were the same as in Figure 2B.

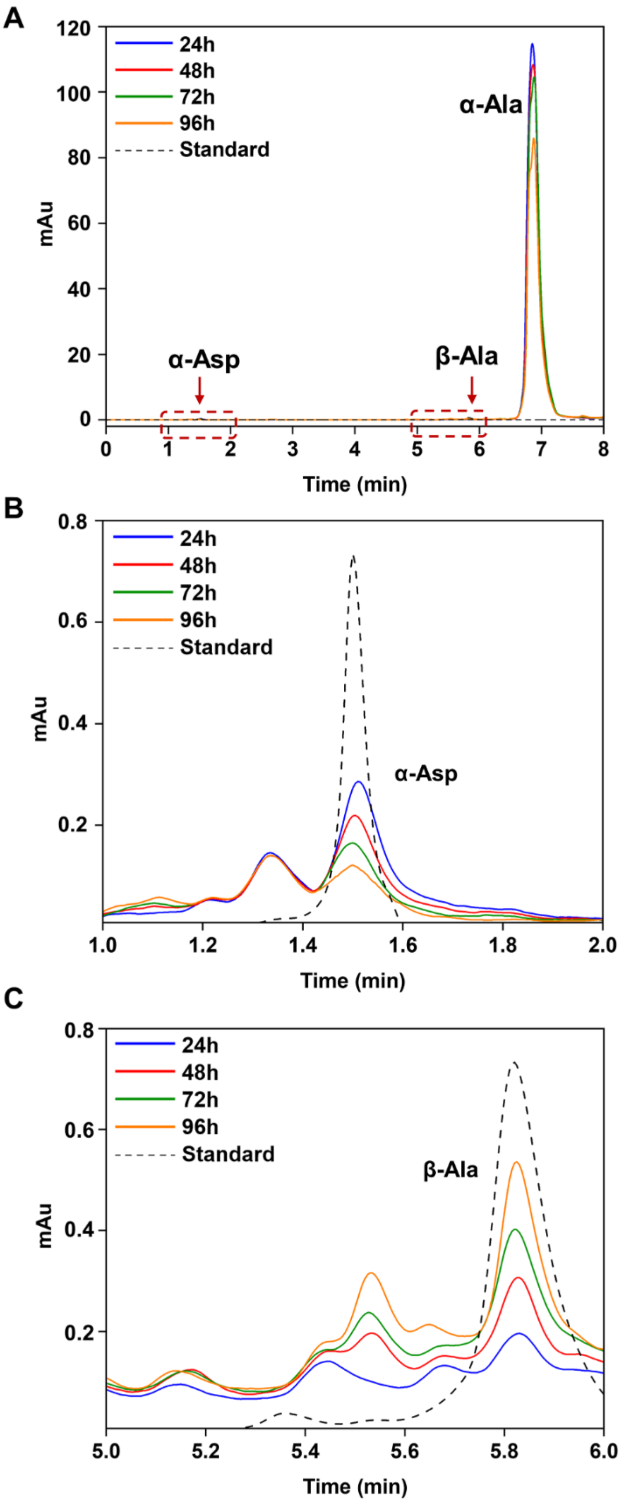

**Figure S14.** HPLC analysis of β-alanine biosynthesis from α-alanine. *In vitro* cascade synthesis of β-alanine at optimal reaction conditions (at 55°C in 100 mM citric acid buffer (pH 4.5), containing 100 mM α-alanine, 1 mM PLP, 100 mM $NaHCO_3$, 10 g/L TsABD, and 0.1 g/L CgAAD). (A) α-Alanine, α-aspartate, and β-alanine in the reaction mixture at 24 h, 48 h, 72 h, and 96 h (solid line) and standards (dashed line). The curves were adjusted to match the retention time for the α-Asp standard peak (B) and β-Ala standard peak (C).

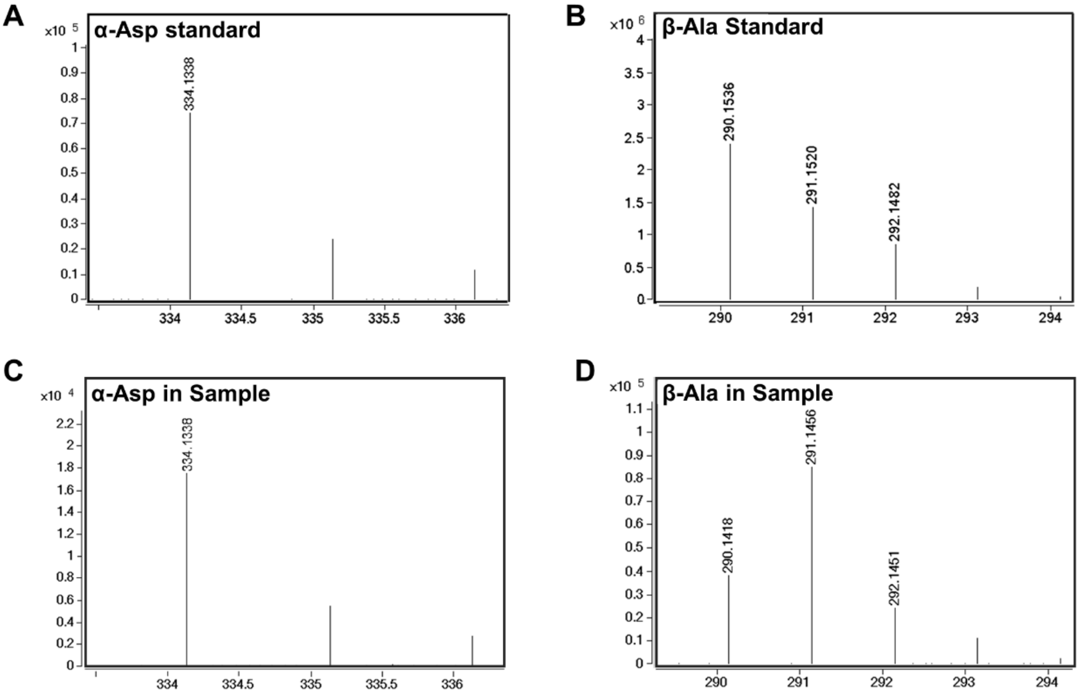

**Figure S15.** Mass spectra of the synthesized α-aspartate and β-alanine. The standard of α-Asp (A) and β-Ala (B) without $^{13}C$-labeling was shown, respectively. (C) The synthesized α-Asp using $^{13}C$-labeling $NaHCO_3$ as the substrate at 96 h. (D) The synthesized β-Ala using $^{13}C$-labeling $NaHCO_3$ as the substrate at 96 h.

### Table Section

**Table S1.** The primers used in this work.

| Primers | Sequence (5' → 3') |
|---|---|
| CgAAD-IF | AGCGGCCTGGTGCCGCGCGGCAGC**CATATG**ctgcgcacCATcctcggaagtaag |
| CgAAD-IR | CAGTGGTGGTGGTGGTGGTG**CTCGAG**ctaaATGcttctcgacgtcaaaagcccggatccaggt |
| CgAAD-VF | cttacttccgaggATGgtgcg**CATATG**ATGGCTGCCGCGCGGCACCAGGCCGCT |
| CgAAD-VR | acctggatccgggcttttgacgtcgagaagCATttag**CTCGAG**CACCACCACCACCACCACTG |
| BsAAD-IF | AGCGGCCTGGTGCCGCGCGGCAGC**CATATG**tatcgaacaATGATGagcggcaaac |
| BsAAD-IR | GCCGGATCTCAGTGGTGGTGGTGGTGGTG**CTCGAG**ctacaaaattgtacgggctggttcg |
| BsAAD-VF | gtttgccgctCATCATtgttcgata**CATATG**GCTGCCGCGCGGCACCAGGCCGCT |
| BsAAD-VR | cgaaccagcccgtacaattttgtag**CTCGAG**CACCACCACCACCACCACTGAGATCCGGC |
| PfAAD-IF | AGCGGCCTGGTGCCGCGCGGCAGC**CATATG**aagttcccaagaaaaggtatcccac |
| PfAAD-IR | CAGTGGTGGTGGTGGTGGTG**CTCGAG**ctaatCATGgcctccgtcaCATGaggCAT |
| PfAAD-VF | gtgggatacctttcttgggaactt**CATATG**GCTGCCGCGCGGCACCAGGCCGCT |
| PfAAD-VR | ATGcctCATGtgacggaggcCATGattag**CTCGAG**CACCACCACCACCACCACTG |
| TkAAD-IF | ATCACAGCAGCGGCCTGGTGCCGCGCGGCAGC**CATATG**tttccagagaggggagcgagcgagg |
| TkAAD-IR | AGCAGCCGGATCTCAGTGGTGGTGGTGGTGGTG**CTCGAG**ttaaagccttttgcaatctccctc |
| TkAAD-VF | cctcgctcgctcccctctctggaaa**CATATG**GCTGCCGCGCGGCACCAGGCCGCTGCTGTGAT |
| TkAAD-VR | gagggagattgcaaaaggctttaa**CTCGAG**CACCACCACCACCACCACTGAGATCCGGCTGCT |
| AfAAD-IF | AGCGGCCTGGTGCCGCGCGGCAGC**CATATG**gcgtggacaacaccctactcgaaatcc |
| AfAAD-IR | GCAGCCGGATCTCAGTGGTGGTGGTGGTGGTG**CTCGAG**ctactgaactttttgatttccctgattttgagc |
| AfAAD-VF | ggatttcgagtagggtgttgtccacgc**CATATG**GCTGCCGCGCGGCACCAGGCCGCT |
| AfAAD-VR | gctcaaaatcagggaaatcaaaaaagttcagtag**CTCGAG**CACCACCACCACCACCACTGAGATCCGGCTGC |
| TtAAD-IF | AGCGGCCTGGTGCCGCGCGGCAGC**CATATG**ttccacgccaagatccaccgggcc |
| TtAAD-IR | GCAGCCGGATCTCAGTGGTGGTGGTGGTGGTG**CTCGAG**ttaccctttgcgcacctcgaggatccg |
| TtAAD-VF | ggcccggtggatcttggcgtggaa**CATATG**GCTGCCGCGCGGCACCAGGCCGCT |
| TtAAD-VR | cggatcctcgaggtgcgcaaagggtaa**CTCGAG**CACCACCACCACCACCACTGAGATCCGGCTGC |

**Table S2.** The plasmids used in this work.

| Plasmids | Description | Source |
|---|---|---|
| TxABD- pET28a | pET28a, contain *aspartate β-decarboxylase* gene from *Thermohalobaculum xanthum* (codon-optimized), $Kan^R$ | This work |
| TmABD- pET28a | pET28a, contain *aspartate β-decarboxylase* gene from *Thermococcus sp.* MV11 (codon-optimized), $Kan^R$ | This work |
| ArABD- pET28a | pET28a carries *aspartate β-decarboxylase* gene from *Acinetobacter radioresistens* DSM 6976 (codon-optimized), $Kan^R$ | This work |
| CtABD- pET28a | pET28a, contain *aspartate β-decarboxylase* gene from *Clostridium thermobutyricum* (codon-optimized), $Kan^R$ | This work |
| TsABD- pET28a | pET28a, contain *aspartate β-decarboxylase* gene from *Thermoactinospora sp* (codon-optimized), $Kan^R$ | This work |
| CgAAD- pET28a | pET28a, contain *aspartate α-decarboxylase* gene from *Corynebacterium glutamicum* ATCC 13032, $Kan^R$ | This work |
| BsAAD- pET28a | pET28a, contain *aspartate α-decarboxylase* gene from *Bacillus subtilis* 168, $Kan^R$ | This work |
| AfAAD- pET28a | pET28a, contain *aspartate α-decarboxylase* gene from *Archaeoglobus fulgidus*, $Kan^R$ | This work |
| PfAAD- pET28a | pET28a, contain *aspartate α-decarboxylase* gene from *Pyrococcus furiosus*, $Kan^R$ | This work |
| TkAAD- pET28a | pET28a, contain *aspartate α-decarboxylase* gene from *Thermococcus kodakarensis*, $Kan^R$ | This work |
| TtAAD- pET28a | pET28a, contain *aspartate α-decarboxylase* gene from *Thermus thermophilus*, $Kan^R$ | This work |

**Table S3.** The strains used in this work.

| Strains | Description | Source |
|---|---|---|
| *E. coli* TOP10 | cloning | Lab stock |
| *E. coli* BL21 (DE3) | expression | Lab stock |
| TxABD-BL21 | pET28a carries *aspartate β-decarboxylase* gene from *Thermohalobaculum xanthum*; BL21 (DE3) | This work |
| TmABD-BL21 | pET28a carries *aspartate β-decarboxylase* gene from *Thermococcus sp.* MV11; BL21 (DE3) | This work |
| ArABD-BL21 | pET28a carries *aspartate β-decarboxylase* gene from *Acinetobacter radioresistens* DSM 6976; BL21 (DE3) | This work |
| CtABD-BL21 | pET28a carries *aspartate β-decarboxylase* gene from *Clostridium thermobutyricum*; BL21 (DE3) | This work |
| TsABD-BL21 | pET28a carries *aspartate β-decarboxylase* gene from *Thermoactinospora sp*; BL21 (DE3) | This work |
| CgAAD-BL21 | pET28a carries *aspartate α-decarboxylase* gene from *Corynebacterium glutamicum* ATCC 13032; BL21 (DE3) | This work |
| BsAAD-BL21 | pET28a carries *aspartate α-decarboxylase* gene from *Bacillus subtilis* 168; BL21 (DE3) | This work |
| AfAAD-BL21 | pET28a carries *aspartate α-decarboxylase* gene from *Archaeoglobus fulgidus*; BL21 (DE3) | This work |
| PfAAD-BL21 | pET28a carries *aspartate α-decarboxylase* gene from *Pyrococcus furiosus*; BL21 (DE3) | This work |
| TkAAD-BL21 | pET28a carries *aspartate α-decarboxylase* gene from *Thermococcus kodakarensis*; BL21 (DE3) | This work |
| TtAAD-BL21 | pET28a carries *aspartate α-decarboxylase* gene from *Thermus thermophilus*; BL21 (DE3) | This work |

**Table S4.** Information of enzymes used in this work.

| Enzyme name | Source organism | Abbreviation | EC number | UniProtKB entry number | Specific activity (U/mg x10$^{-3}$) | Conditions |
|---|---|---|---|---|---|---|
| aspartate β-decarboxylase (α-Ala→α-Asp) | *Thermohalobaculum xanthum* | TxABD | 4.1.1.12 | *A0A8J7M890* | 0.14 | pH 4.5, 37°C |
| | *Thermococcus sp. MV11* | TmABD | 4.1.1.12 | *A0A8J8A898* | 0.12 | pH 4.5, 37°C |
| | *Acinetobacter radioresistens DSM 6976* | ArABD | 4.1.1.12 | NCBI: *BBL20616.1* | 0.10 | pH 4.5, 37°C |
| | *Clostridium thermobutyricum* | CtABD | 4.1.1.12 | *N9Y2J5* | 0.11 | pH 4.5, 37°C |
| | *Thermoactinospora sp* | TsABD | 4.1.1.12 | *A0A8J8HAW7* | 0.20<br>1.35 | pH 4.5, 37°C<br>pH 3.5, 55°C |
| aspartate α-decarboxylase (α-Asp→β-Ala) | *Corynebacterium glutamicum ATCC 13032* | CgAAD | 4.1.1.11 | *Q9X4N0* | 10.10 | pH 7.5, 37°C |
| | *Bacillus subtilis 168* | BsAAD | 4.1.1.11 | *P52999* | 5.40 | pH 7.5, 37°C |
| | *Archaeoglobus fulgidus* | AfAAD | 4.1.1.11 | *A0A075WHK3* | 1.29 | pH 7.5, 37°C |
| | *Pyrococcus furiosus* | PfAAD | 4.1.1.11 | *Q8U1P6* | 0.34 | pH 7.5, 37°C |
| | *Thermococcus kodakarensis* | TkAAD | 4.1.1.11 | *Q5JJ82* | 0.80 | pH 7.5, 37°C |
| | *Thermus thermophilus* | TtAAD | 4.1.1.11 | *Q72L22* | 0.17 | pH 7.5, 37°C |

Results are means ± SD of three replicates.

**Table S5.** List of sequences of the amplified or synthesized genes.

| Gene | Sequence |
|---|---|
| *TxABD* | atggatcgcagctattttgagagcctgagcgtgctgagcccgtttgaaattaaagatgaactgatta<br>aactggcgcgcgcgggcagcgaacgcagcgcgaaagcgtttctgaacgcgggccgcggcaa<br>cccgaactggattgcgagccgcgcgcgcgaagcgtttttctgctgggtcagtttgcgctgattgaa<br>gcgaaacgcacctatcatgatccggaagcgggcattgcgggcatgccgccgcgcgatggcgcg<br>tatattcgctttagccgctgggtgaaacgtcagcaacaagcgcaagaagaaggctggattgaag<br>aagatggcgtgccgccggcgcgcaccctgggcgatgcggtggattttgcgattgaaaaatttggct<br>ttgatccggatgattttgtgcatgaactgaccgatagcattattggcgataactatccggtgccggat<br>cgcgcgctgcatcataacgaaattattacccatgaatatctgatgtgggcgatgtgcgcgggccat<br>cgcccggatgtgaaatttgatctgtatccggtggaaggcggcaccgcggcgatgtgctatattttta<br>aagcgctgaaaaatcagcgcctgctgaacccgggcgataccattgcgctgggcaccccgattttt<br>accccgtatctggaaatgccggaactggaagattatgcgctgcataccattaacgtgagcgcgga<br>acaagaagataaatttcagctgaccgaagatgatctgaaagcgctggaagatccgaaagtgaa<br>agcgctgtttctggtgaacccgggcaacccgagcgcggccgcgatggatccggatgcgattgcg<br>cgcctggtgaacctggtgaacaccaaacgcccggatctgatgattctgaccgatgatgtgtatggc<br>acctttgtgccgggctttgaaagcctgttaggccatctgccgcgcaacaccattggcgtgtatagct<br>atagtaaatattttggctgcaccggctggcgcctgggcgtgattgcggtggcgcaagataacattttt<br>gatacccgcattcaagaacatccggaagaaattcagaaactgctggatgatcgctatcgcgcgct<br>gaccccgaaaccgcgcgaaattaaatttattgatcgcgtggtggcggatagccgcgatgtggcgc<br>tgaaccatacgagcggcctgagcctgccgcagcaagtgatgatgaccctgtttggcctggtggaa<br>atgatggatagcgaaaaagtgtatcagaaagcgtgcattgcgatttgcgaacgccgctttcgcaa<br>cctggtggaaggcatgggcgtggaaagcaacccgggcaaatattttgattattactatggcattatt<br>gatctggaattttggctgcgcaaatatgtgggcgaagaagtggtgcagtgggtgaaagaaaacat<br>tcatccgctggatattccgtttcgcctggcggaagatcatggcattgtgctgctgaacggcagcggc<br>tttgcggcgccgaactggagcgcgcgcgtgagctttgcgaacctgcatgatgatgcgtatagcga<br>aattggccgcgcggtgcgcagcattgcgcgcggctatgtgcaagcgtatcaagcgagccgcgg<br>cgaagaaattaccgcgtaa |
| *TmABD* | atgctgaaaaacattggcaacctggaagaactgagcccgtttgaatttaaagaactgctgattaaa<br>ctggcgcgccgcaaaagcgaacgcatgatgctgaacgcgggccgcggcaacccgaactttctg<br>gcgctgaccccgcgctatgcgtatctgcagctgggcaaatttgcgctgagcgaagcggaacgcc<br>attttggctatatgggcggcctgattggcggccatagcgatcgcgaaggcattgaagcgcgctttg<br>aaatttttgtgcgcaaccattggaacgaacgcggcaccgcgtttctgaacagcgcggtgagctatg<br>tgcgcgattatctgggcctgagcgcgggcgattttctgcatgaaatggtgcaaggctatctgggctg<br>cgattatccgagcccgccgcgcatgctgccgctggcggaaaaaattgtggcgcgctatctgatga<br>aagaaatgggcgcgggctatgatctgggctatagcctggtggatgaaacgcagctgtttgcggtg<br>gaaggcggcaccgcggcgatggcgtatctgtttgaaagcctgaaagcgaaccgcattctgaacg<br>aaggcgataaaattgcgctggcggtgccgatttttagcccgtatctggaaattccgcgcctggatac<br>ctatcgcctggaagtgattgaagtgcgcgcggatgaagaacgcggctatcagattccgcgcgaa<br>gaactggaaaaactgcgcgatccggaaattaaagcgtttttctggtgaacccgggcaacccgac<br>gagcgtgaaactggaagaaggcaccctggaagatctgcgcgaaattgtggaaaaagatcgca<br>acgatatgattatcattaccgatgatgtgtatgcgacctttgcggaagattttcgcagcgtgtatagcg |

| | |
|---|---|
| | tgctgccgcataacaccattctggtgtatagctttagcaaatattttggcgcgaccggctggcgcctg<br>ggcgtgattgcgctgcatcgcgataacgtggtggatcgcctgattgcgagcctgccggaagaagt<br>gcaagaaattctggaaaaacgctatgcgccgattaccccggatgtgcgcgaactgaaatttattg<br>atcgcttagtggccgacagccgcaacgtggccctgcgccatacggcgggcctgagcaccccgc<br>agcaagtgcagatggtgctgtttgcgctgtatgcgctgatggatgaagaggaacgctataaaaaa<br>accgtgaaacatgtgctgcgccgtcgctatcgcgcgctgtatcgcggcattggcattgaaccgga<br>agaaagcccgagctatgcgtattactataccctgctggataccgaaaaactggcggaacgcctgt<br>atggcaaagaatttgcggaatggtttgtgcgcaccctgccggtggaagaatttattgtgcgcctggc<br>ggtggaagcgggcgtggtgctgctgccgggcaaaggctttgatgtggtgcatccgagcgcgcgc<br>gtgagcctggcgaacctgcgcgagattgactatattaaaattggcaaaaccattcgccgcctgatt<br>gatgaatattataaaaaatttcgcggcgaagtgtaa |
| *ArABD* | atgggcaacgtggattatagcaaatatagcaaactgagcccgtttgaactgaaagatagcctgatt<br>gcgctggcgcagagcaaacgcgatcgcctgatgctgaacgcgggccgcggcaacccgaacttt<br>ctggcgaccctgccgcgccgcgcgtttttcagctgggcttatttagcgcgaccgaaagcgaattta<br>gctttagctatatgccggaaggcctgggcggctttccgcgcccggtgggcctgcagagccgctttg<br>ataactttctgatgcagaaccgcgataaaccgggcgtgctgtttctgggcaaagcggtgagctatg<br>tgcgcgatcagctgggcctggacccggatatgtttctgctggaaatggtggaaggcattctgggct<br>gcaactatccggtgccggatcgcatgctgcgcgtgagcgaaaccattattaaagaatatctgctgc<br>aagaaatgggcattaaaagcatgccgaaagaaggcctggatctgtttgcggtggaaggcggca<br>ccgcggcgatggcgtatatttttaacagcctgaaagaaaaccgcattattaacaccggcgatcgc<br>attgcgattggccgcccgatttttaccccgtatctggaaattccgaaactgaacgattatcagctgga<br>agaaatttttattgaagcggatccgaacctgggctggcagtatccggaaagcgaactgcgcaaa<br>ctggaagatccgagcattaaagcgtttttctggtgaacccgagcaacccgccgagcgtgaaaatt<br>agcgatgaaggcctgctgattctggcggatattgtgcgcaaacgcccggatctgattattctgaccg<br>atgatgtgtatggcacctttgcggatgattttaaaagcctgtttgcgatttgcccgaacaacaccattct<br>ggtgtatagctttagcaaatattttggcgcgaccggctggcgcctgggcattattgcgctgagcaac<br>aataacattattgatcagaaaattgcggcgctgagcgatcaagaaaaacaagaactggaagaa<br>cgctatagcagcctgaccaccgaaccggaaaaaattaaatttattgatcgcctggtggcggatag<br>ccgcaacgtggcgctgaaccataccgcgggcctgagcaccccgcagcaagtgcagatggtgct<br>gtttgcgctgtttaacatgatggatagccgccaagcgtataaaaaagcggtgaaaagcgtggtgc<br>gcgaacgcgatgcggcgctgtatcgtcagctgggcgtggaagtgccggaagatctgaacgcggt<br>ggattattataccctggtggatctggaacgcaccgcgcgcattctgtatggcgatgattttgcgaact<br>gggtgatggtgaacaaaaacccgaccgaactgctgtttcgcgtggcggatgaaaccggcgtggt<br>tctgctgccgggtagcggcttcggcgttagccatccgagcgcccgcgcgagcctggcgaacctg<br>aacgcgtatcagtatgcggcgattggcgatagcctgcgccgctttgcggaagatgcgtatcaaga<br>atatctgggcaccaaaaaagatgaaagctaa |
| *CtABD* | atggatcgcaacattcagcgcgaagaaattgaaaaaatttatggcaaaattagcccgtttgaattta<br>aaaacaaactgattagtctggcgacggaggaaaagggcgcccgcaccctgctggatgcgggtc<br>gcggcaacccaaactggacctgcgcgaccccgcgccaagcgtttttttacctttggtcagtttgcggt<br>ggaagaaacgcagcgcacctggcagaactgggatctggcgggcatgccgagcaaaagcggc<br>atttatgaacgctttctgaactatattaaagaaaacccgaacatgccgggcattgaactgctgaaa<br>gatatgattgaatatggcattaacaacaaaggctttaccccggatgattgggtgtttgaactggcgg<br>atggcattattggcgataactatccgtttccggatcgcatgctgaaccatattgaacagattgtgcat<br>gattatctgattcaagaaatgaaatatcgcaacgaaaaagatgataaatttaacatttatgcggtgg |

| | |
|---|---|
| | aaggcgcgaccgcggcgatgtgctatctgtttgatagcctgaaagtgaaccatctgctgaacgaa<br>ggcgataccattgcgattatgaccccgatttttaccccgtatctggaaattccgattctgccgaactat<br>aacctgaacgtggtgaaaattcaagcgagcgaagaaagcacgagcggcgtgcatacctggca<br>gtatccgaaagaagaactggataaactgaaagatccgaacattaaagcgctgtttattgtgaacc<br>cgagcaacccgccgagcattgcgctgagcgatcgctgccgcaaccatattgtgagcattgtgcaa<br>gaatataacccgaacctgattatcattagcgatgatgtgtatggcacctttgtggaaggctttcgcag<br>cctgatgaacgatctgccgtataacaccattggcgtgtatagctttagcaaatattttggcgtgaccg<br>gctggcgcctgggcaccattgcgctgcatgaaaaaaacattctggataaattaattcgcgaactgc<br>cgagcgaacataaagaaaccctgaaccgccgctatggcgatatgtgcaaagatccgagcacc<br>gtgccgtttattgatcgcattgtggcggatagccgccaagtggcgctgaaccataccgcgggcctg<br>agcaccccgcagcaagtgcagatggcgttttttgcatgtttgcgctgctggataaagcgaaccgct<br>ataaaaacaaaaccattgatatttgccatcatcgtcagaaactgctgtttgaaggcctggatctgga<br>actgccgaaaaacaaatatgatgcggcgtattatacgcagtttgatctgctggaatgggcgaaca<br>aatattatggcagcgaatttgcggcgtatctgcagaaatatcataaaccgagccatattctgtatcg<br>cctggcgcaagaaagcagcattgtgctgctgagcggcagcggctttcaaggcccggaatggag<br>cattcgcattagcctggcgaacctgaacgataacgcgtatagcaaaattggcaacgtgctgcata<br>acattctggaagaatttgtgaaagcgtgggaagaaaccaaacagcagtaa |
| *TsABD* | atggcggatcgcgcgaccgaaagcaaatgggaagcgctgagcccgtttgaactgaaaaacga<br>actgattgatctggcgaaagataacaaagcggcgcataccatgctgaacgcgggccgcggcaa<br>cccgaactggattgcgaccgaaccgcgcgaagcgttttttctgctgggcagctttggcctggcgga<br>aagccgccgcacctggagcgaatttgatggcctgggcggcatgccggcgcaagaaggcattgc<br>gggccgctttggcgcgtttctggatgatcatcgcgatcgcccgggcgcggaactgctgcgcaaaa<br>gcgtggaatatggcacgagcaccctgggctttgaagcggatgcgtttgtgtgggaactggcggat<br>gcgattattggcgatcattatccggaaccggatcgcatgctgcatcatgcggaacgcattgtgcatg<br>cgtatctggtgcaagaaatgtgcgcgggccaagcgccggcgcgcccgtatgatctgtttgcggtg<br>gaaggcggcaccgcggcgatgtgctatatttttgatacctgcgtgattaacggcattctgcgcccgg<br>aagataccattgcgattatgaccccgatttttaccccgtatctggaaattccgcatctggaacgctat<br>cagtttaaaaccgtggaaattgcggcgagcaaaatgaaccaagatggcctgcatacctggcagt<br>atgatcaagcggaactggataaactggcggatccgaacgtgaaagcgctgtttctggtgaaccc<br>gagcaacccgccgagcgtgatgctggcgcaagaaagcctggatcatctggaaaaaattattgcg<br>acgagcaacccggatctgaccattattaccgatgatgtgtatggcacctttgtgaacggctttcgca<br>gcctgatgagcaccgcgccgcgcaacaccattggcgtgtatagctttagcaaatattttggctgca<br>ccggctggcgcctgggcgtgattgcggtgaaccaagataacatttttgaaggcaaactggcgggc<br>ctgccgcagagcgataaagatcgcctgagcgatcgctatagcaccattagcctggatccggcgg<br>gcattacctttattgatcgcctggtggcggatagccgccaagtggcgctgaaccataccgcgggcc<br>tgagcctgccgcagcaagtgcagatgagcctgtttagcctggcggcgctgctggataccgatgat<br>acctataaaaaaacctgccaagcgattattgcgcgccgcctgaaagcgctgaccgaaggcatg<br>ggcgtgccgctgccggatgatccgctggcggcgcgctattatgtggaactggatattctgaactatg<br>tggaaaaaaacctggggcaaagaatttgcgggctggctggaacagaactttgaaccggtggatcc<br>ggtgtttcgtttagcggaaaaaggtagcgtggtgttactgaacggtggcggttttgatggcccgggtt<br>ggagcgtgcgcgtgagcctggcgaacctggcgcaagatgattatgcgcagattggcacctggct<br>gcgcgaagtggtggaaggctataaagcggaatttgataaaagctaa |
| *CgAAD* | atgctgcgcaccatcctcggaagtaagattcaccgagccactgtcactcaagctgatctagattat<br>gttggctctgtaaccatcgacgccgacctggttcacgccgccggattgatcgaaggcgaaaaagt |

| | |
|---|---|
| | tgccatcgtagacatcaccaacggcgctcgtctggaaacttatgtcattgtgggcgacgccggaa<br>cgggcaatatttgcatcaatggtgccgctgcacaccttattaatcctggcgatcttgtgatcatcatga<br>gctaccttcaggcaactgatgcggaagccaaggcgtatgagccaaagattgtgcacgtggacgc<br>cgacaaccgcatcgttgcgctcggcaacgatcttgcggaagcactacctggatccgggcttttgac<br>gtcgagaagcatttag |
| *BsAAD* | atgtatcgaacaatgatgagcggcaaacttcacagggcaactgttacggaagcaaacctgaact<br>atgtgggaagcattacaattgatgaagatctcattgatgctgtgggaatgcttcctaatgaaaaagt<br>acaaattgtgaataataataatggagcacgtcttgaaacgtatattattcctggtaaacggggaag<br>cggcgtcatatgcttaaacggtgcagccgcacgccttgtgcaggaaggagataaggtcattattat<br>ttcctacaaaatgatgtctgatcaagaagcggcaagccatgagccgaaagtggctgttctgaatg<br>atcaaaacaaaattgaacaaatgctggggaacgaaccagcccgtacaattttgtag |
| *AfAAD* | atggcgtggacaacaccctactcgaaatccggaaggtcggctctcgtgcctgctgggccgtgga<br>gctacggtatggaggccatatccgtccatgcgatgcttagcggctttgaggagttgctgccggaaa<br>acatggaaagtgagggggaggggtatgtttactttgccgaaatcatctcgcaaagtccgaactttc<br>ccgagctcaactacgaagctccgggactggttcagtacagagagcttgccatattcctgaaggtg<br>aaatatggcgggagagtcctcgcctactgcccctttatgtacgttgacaacgatgtctcactgctga<br>gaggctttgttgccggatttccaaaaaagcttgctgtcattgacattacaagggagcatcctctgcttg<br>agcagaaaaaattcgggggagttgcgatgaggggcggttacggccttgcaaagatggtggttga<br>gccgatggaggaggtcgagtccaatccgcttgacagctttggaagctggatgctctacaggtttgc<br>cgaacctatgggtgtaaaggagttcgtggagatagttcctgaggttagctacgcgaaaatcgaga<br>gaggaaacgcagagctgaagctttacagtgggataaacgacgagttgcctgttgtaaaggaaat<br>tcttggggggttacaggttctcggccttgctcaaaatcagggaaatcaaaaaagttcagtag |
| *PfAAD* | atgaagttcccaagaaaaggtatcccacaagaggaagttatgagagaattagaaaaatacactt<br>ccaaagacttatcattttcatctggcaagatcctaggttccatgtgcactctcccgcacgagttggcc<br>aaagaagtgttctgtatgtacatggatagaaatttgggcgatcctgggcttcacccagggacaaag<br>aaaatagaagaagaagtaatagaaatgctttcagatctcttacatttagagagaggatatggacat<br>attgtctcaggtgggactgaggcaaatatcttggctgtcagagcgtttagaaatctggcagatgttga<br>aaatccagagttaattttgcctaagagtgcacacttctccttcataaaggctggagaaatgttggga<br>gttaagttgatttgggctgacttaaatccagactatactgtggatgtaaaagatgttgaggcaaagat<br>aagtgagaataccattggcatagttggaatagccggaactacagggcttggtgtcgttgacgatatt<br>ccagctttgagtgatctagctagggactatggaatccctcttcacgttgacgcagcatttggaggattt<br>gtaataccatttgctaaagaactcggatatgacttaccagactttgacttcaagcttaaaggagttca<br>aagtataacaatagatccccacaagatgggaatggccccaattccggcaggggggaatagttttta<br>gacataaaaaaatatctaagggcaataagcgttttagctccatatttagctggagggaagatatggc<br>aggcgacaataacgggaactagaccaggagctagtgtgttggctgtatgggctctaattaagcatt<br>taggatttgaagggtacatggagattgtagatagggccatgaagctttctaggtggtttgctgagga<br>gataaagaaaactccaggagcttggctggtgagagagcccatgctaaacatagtctcctttaaga<br>ccaagaacctaaggagagttgaaagggaattaaaatcaaggggatggggaattagtgcccata<br>gggggtatataagaatagtttctcatgcctcatgtgacggaggccatgattag |
| *TkAAD* | atgtttccagagaggggagcgagcgaggaagaagtcctgagggagcttgaagagaagacaa<br>gagaagatttaaccttcgattcaggcaagatactcggttctatgtgcacttacccacaccccttcgca<br>gtgaaggtcgttatgaagtacatagacaggaacctcggcgatcctggtttacatatcgggagcca<br>gaagatcgagaaagaagccgttgacatgctggccaacctacttgggctggagaagggctacgg<br>ccacatagtctctggcgggacggaggccaacattctggcagtgagggcaatgagaaacctcgct |

| | |
|---|---|
| | ggaatagaaaagcccgaactgatccttccagagagcgctcacttctcattcataaaggccgccg<br>agatgctaggggtaaagctcgtctgggcggaactgaacgacgactacacagtcaacgtgaaag<br>acgttgagaagaagataacagacaggacgataggaatagtggggatagctggaacgaccgg<br>ccttggagtcgtcgacgacatcccggccctgagcgacttagcacttgactacggtcttcccctccac<br>gttgatgcagccttcggcggcttcgtgataccctttgcaaaggctctcggttatgagattcccgacttc<br>gacttcaggctcaagggagtgaaaagtataacaatagaccctcacaagatggggatggtaccg<br>attccagccgggggaataatcttccgcgagaagaagtttctggacagtataagcgttttagccccgt<br>atttggccggaggcaaaatctggcaggcgacgataaccggaaccagaccgggagccaatgct<br>ctggcagtgtgggcgatgataaaacacctcggcttcgatgggtacaaggaagttgtaaaagaga<br>agatggagctcgcaaggtggttcgcctcggaactgaagaagattccagggatttacctcatcagg<br>gagccagtgttaaatatagtctcattcggttcagagaagctggaagagcttgaaaaagagcttaa<br>ggcgaggggctggggtgtaagcgctcacaggggctacataaggatcgtcgtcatgcctcacgtg<br>aagagggagcacctcgaggagtttttgagggatttgagggagattgcaaaaaggctttaa |
| *TtAAD* | atgttccacgccaagatccaccgggccacggtgacccaggccgacctccactacgtgggctcgg<br>tgacggtggaccaggacctcctggacgccgccgggatccttccttttgagcaggtggacatctacg<br>acatcaccaacggggcccggctcaccacctacgcccttcccggggaacgggggtccggcgtca<br>tcgggatcaacggggccgcggcccacctggtgaagcccggggacctggtcatcctcgtggccta<br>cggcgtctttgacgaggaggaggcgaggaacctcaagcccaccgtggtcctggtggacgagag<br>gaaccggatcctcgaggtgcgcaaagggtaa |
